# Mitochondrial STAT3-mediated suppression of apoptosis constrains antimycobacterial immunity

**DOI:** 10.64898/2026.06.26.734825

**Authors:** CJ Mabry, AK Coleman, MH Smith, SL Hahn, TM Newbolt, MJ Chapman, LW Stranahan, CG Weindel, KL Patrick, RO Watson

**Author notes:** Address Correspondence to Robert O. Watson.

## Abstract

Maintenance of mitochondrial homeostasis is required to balance the host-pathogen interface during *Mycobacterium tuberculosis* (Mtb) infection. Here, we identify the non-canonical TRIM family member Trim14 as a critical regulator of mitochondrial integrity in Mtb-infected macrophages. Specifically, we demonstrate that Trim14 preserves mitochondrial membrane polarization and limits macrophage apoptosis by controlling phosphorylation and mitochondrial targeting of Stat3. When targeted to mitochondria, Stat3 restricts opening of the mitochondrial permeability transition pore, which raises the macrophage threshold for apoptotic commitment. *In vivo*, loss of Trim14 enhances apoptosis of macrophages and dendritic cells, leading to augmented antimycobacterial immunity marked by increased CD8^+^ T cell activation and effector function. Together, these findings define a Trim14–mitochondrial Stat3 axis that suppresses host-protective apoptosis during Mtb infection and pinpoint mitochondrial Stat3 as a potential target for therapies aimed at boosting antimycobacterial immunity.

**HIGHLIGHTS:**

- Trim14 raises the apoptotic threshold in Mtb-infected macrophages.
- Trim14 controls phosphorylation and mitochondrial targeting of Stat3.
- Reduced mitochondrial Stat3 promotes mPTP opening and apoptotic commitment.
- Trim14 deficiency enhances apoptosis, CD8^+^ T cell immunity, and Mtb resistance.

## INTRODUCTION

During *Mycobacterium tuberculosis* (Mtb) infection, cell death pathways dictate whether infected cells restrict bacterial growth or promote dissemination^1^. Apoptosis, a form of programmed cell death characterized by maintenance of membrane integrity, is largely protective during Mtb infection, as containment of bacilli within apoptotic bodies limits bacterial spread and reduces tissue damage^2–5^. Apoptotic cell death further enhances antimycobacterial immunity by promoting efferocytosis, antigen cross-presentation, and protective T cell responses^6–10^. Inflammatory forms of cell death, on the other hand, such as necroptosis, inflammasome-mediated pyroptosis, and ferroptosis, are associated with worsened Mtb outcomes^11–16^. These lytic cell death modalities can compromise infected macrophages, release bacilli into the extracellular space, amplify inflammatory cytokine production (e.g. type I IFN), and worsen lung tissue pathology^17–19^. Consistent with cell death playing a key role in dictating Mtb outcomes, Mtb has evolved ways to polarize macrophages to push them towards pro-bacterial cell death modalities. For example, the ESX-1 secretion system promotes phagosomal membrane damage and cytosolic access of Mtb virulence factors, triggering inflammatory macrophage death and facilitating bacterial spread^20,21^. Likewise, virulent Mtb strains inhibit apoptosis by expressing proteins such as NuoG and SecA2 that limit mitochondrial apoptotic signaling and oxidative stress-induced apoptosis^22,23^. Thus, the balance between apoptosis and inflammatory cell death represents a critical determinant of host resistance to tuberculosis infection.

Mitochondria serve as central metabolic and signaling platforms that coordinate innate immune responses in Mtb-infected macrophages^24–26^. During infection, macrophages undergo extensive bioenergetic remodeling, dynamically shifting how they make and store energy downstream of pathogen sensing and other inflammatory cues. In addition to controlling cellular bioenergetics, mitochondrial fitness—defined by membrane potential, respiratory capacity, organelle integrity, and quality-control pathways— is a key determinant in driving cell death modality usage^24,27–30^. We know that mutations in host mitochondrial genes are associated with chronic inflammation and increased susceptibility to mycobacterial disease^27,29,31,32^. We also know that Mtb perturbs mitochondrial homeostasis through ESX-1-dependent phagosomal membrane damage, which exposes the host cytosol to Mtb virulence factors that promote mitochondrial stress and necrosis^20,24,25^. However, despite clear links between mitochondrial dysfunction and Mtb pathogenesis, it has been difficult to identify mitochondrial-associated proteins that play clear pro- or anti-bacterial roles in Mtb-infected macrophages or in pre-clinical models of tuberculosis.

Members of the multifunctional tripartite motif-containing (TRIM) family of innate immune regulatory proteins are well-positioned to regulate the host-pathogen interface. Canonical TRIM proteins are defined by a conserved RBCC architecture composed of RING, B-box, and coiled-coil domains, with the RING domain often conferring E3 ubiquitin ligase activity^33^. A smaller number of TRIMs, among them TRIM14, comprise a non-canonical subset of TRIM proteins that lack a RING domain and function via protein–protein interactions through their C-terminal PRY-SPRY domain^34^. Notably, beyond its role in innate immunity^35–37^, TRIM14 has been linked to cell survival programs in cancer. Elevated TRIM14 expression is associated with poor outcomes in several malignancies, including colorectal, hepatocellular, papillary thyroid, and breast cancers^38–41^. Mechanistically, TRIM14 has been reported to promote tumor progression by restraining apoptosis and modulating autophagy, proliferation, and inflammatory signaling pathways^38,39,42,43^. These findings position TRIM14 as a regulator of both immune signaling and cell fate across various biological contexts.

We previously found that Trim14 scaffolds Tbk1 and Stat3 during Mtb infection of murine macrophage cell lines, promoting Stat3 activation and limiting excessive type I IFN responses^37^. Signal Transducer and Activator of Transcription 3 (STAT3) is best known as a transcriptional regulator of inflammatory and pro-survival programs in cancer and infection, where it can promote resistance to apoptosis and shape immune cell function^44–47^. More recently, we have come to appreciate non-canonical roles for STAT3 at mitochondria, where it promotes electron transport chain activity and reduces sensitivity to oxidative stress^48–50^. Here, we identify Trim14 as a regulator of mitochondrial homeostasis that sets the threshold for apoptotic cell death in Mtb-infected macrophages. We report that macrophages lacking Trim14 exhibit increased sensitivity to intrinsic apoptosis triggers and impaired mitochondrial function, due to defects in phosphorylation and mitochondrial targeting of Stat3. *In vivo*, loss of Trim14 elicits host protective mechanisms—namely increased CD8+ T cell activation and effector functions—resulting in enhanced survival of Mtb-infected mice. Our data highlight an unappreciated role for mitochondrial Stat3 in promoting apoptosis of Mtb-infected macrophages and argue that therapeutic targeting of the Trim14-Stat3 axis could help boost antimycobacterial immunity in patients.

## RESULTS

### Trim14 limits cell death in response to apoptotic triggers and Mtb infection

Previous work from our lab has established a role for Trim14 in regulating type I IFN expression during Mtb infection of a murine macrophage cell line (RAW 264.7)^37^. Underlying this phenotype, we found that Trim14 interacts with the innate immune kinase Tbk1 and influences Tbk1’s ability to phosphorylate substrates, including the transcription factor and non-canonical regulator of mitochondrial health, Stat3. Because TBK1 and STAT3 have previous links to cell death and because balancing cell death modalities is critical for controlling Mtb pathogenesis, we asked how Trim14 affects the propensity of primary macrophages to undergo different types of cell death. Briefly, we isolated bone marrow derived macrophages (BMDMs) from WT and *Trim14^-/-^* mice (**Fig. S1A-B**), infected them with Mtb (Erdman, MOI=5), and followed propidium iodide (PI) incorporation over time. We did not observe any spontaneous cell death in *Trim14^-/-^*BMDMs (**Fig. S1C**), but Mtb triggered more cell death in *Trim14^-/-^* BMDMs relative to WT controls over a 24h infection (MOI=5) (**Fig. 1A**). Given that Mtb infection of macrophages simultaneously elicits several forms of cell death (e.g., apoptosis ^2,3^, necroptosis ^5^, and inflammasome-mediated pyroptosis^13,15^), we used a panel of standard agonists to identify the cell death modalities at play. We measured no Trim14-mediated differences in cell death triggered by the NLRP3 inflammasome (3h LPS + ATP) (**Fig. S1D**), the AIM2 inflammasome (poly dA:dT) (**Fig. S1E**), or extrinsic necroptosis (TNFα/birinapant/z-VAD-FMK) (**Fig. S1F**). We did, however, observe dramatic differences in cell death when *Trim14^-/-^* BMDMs were treated with apoptotic triggers including etoposide, which induces DNA breaks (**Fig. 1B**), the kinase inhibitor staurosporine (**Fig. 1C**), or the BCL-2 inhibitor ABT-737 (**Fig. 1D**). Enhanced cell death in response to Mtb, staurosporine, and ABT-737 was also observed in *Trim14^-/-^* BMDMs immortalized via the Cre-J2 retrovirus (iBMDMs; **Fig. 1E, S1G, 1F**), providing us a more genetically tractable system for studying the mechanisms underlying these phenotypes.

**Figure 1.**
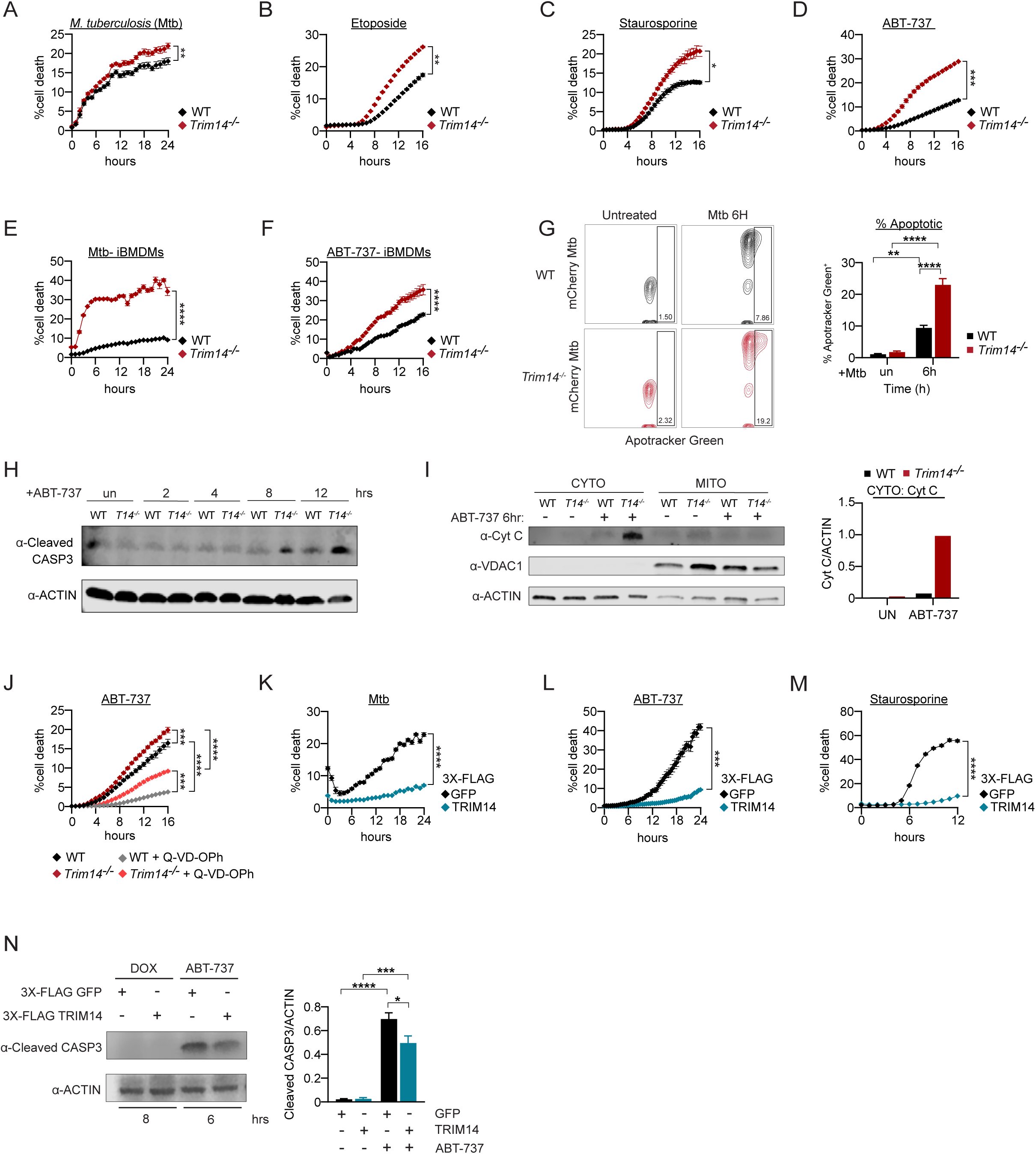
Trim14 is a negative regulator of apoptotic cell death in macrophages. A) Cell death in WT and *Trim14^-/-^* BMDMs infected with WT Mtb (MOI=5). Cell death was measured by propidium iodide uptake (% cell death = propidium iodide positive/total cell count × 100%). B) As in (A) but treatment with 25 µM etoposide. C) As in (A) but treatment with 0.1 µM staurosporine. D) As in (A) but treatment with 10 µM ABT-737. E) As in (A) but Mtb infection (MOI=5) in WT and *Trim14^-/-^* immortalized BMDMs. F) As in (E) but treatment with 10 µM ABT-737. G) Flow cytometry analysis of apotracker green in WT and *Trim14^-/-^* iBMDMs infected with WT Mtb (MOI=5) for 6hr. Quantification of % apoptotic (% apotracker green positive) H) Immunoblot of cleaved caspase 3 in WT and *Trim14^-/-^* iBMDMs treated with 10 µM ABT-737 (2, 4, 8, 12hr). Actin was used as a loading control. I) Cytochrome C release measured by biochemical fractionation and immunoblot analysis in untreated and ABT-737 treated (10 µM 6hr) WT and *Trim14^-/-^* iBMDMs. Vdac1 was probed to control for mitochondria enrichment and mitochondria contamination in the cytosol. Quantification on right shows Cytochrome C relative to Actin. J) Cell death in WT and *Trim14^-/-^* iBMDMs treated with 10 µM ABT-737 and 20 µM Q-VD-OPh. 20 µM Q-VD-OPh was preincubated 1hr before addition of ABT-737. | K) As in (A) but in tetracycline inducible overexpression RAW 264.7 macrophages (treated with 333ng/ml doxycycline for 4hr pre-infection) expressing 3X-Flag GFP or 3X-Flag TRIM14. L) As in (K) but treatment with 10 µM ABT-737. M) As in (K) but treatment with 0.1 µM staurosporine. N) Immunoblot of cleaved caspase 3 in tetracycline-inducible overexpression iBMDMs expressing 3X-Flag GFP or 3X-Flag TRIM14. After 8hr doxycycline pre-treatment, cells were treated with 10 µM ABT-737 for 6hr. Quantification of cleaved caspase 3 normalized to Actin on right. Statistical analysis: *p < 0.05, **p < 0.01, ***p < 0.001, ****p < 0.0001. Statistical differences were determined for (A-G, J-N) using two-way ANOVA with Tukey’s post-test.

Initiation of apoptosis results in the translocation of phosphatidylserine to the outer plasma membrane, which can be detected via flow cytometry using the fixable dye Apotracker Green or Annexin V. Consistent with increased PI incorporation in *Trim14^-/-^* macrophages, we measured increased Apotracker green on the surface of *Trim14^-/-^* iBMDMs at 6h post-Mtb infection (**Fig. 1G**) and increased annexin V following ABT-737 stimulation (**Fig. S1H**), as well as enhanced caspase-3 cleavage via western blot (**Fig. 1H**). We also measured an increase in cytochrome c release into the cytosol of *Trim14^-/-^* macrophages following ABT-737 treatment via cellular fractionation (**Fig. 1I**). Importantly, we were able to rescue most cell death in *Trim14^-/-^* BMDMs with the pan-caspase inhibitor Q-VD-OPH, indicating that enhanced cell death in *Trim14^-/-^* cells is caspase-dependent (**Fig. 1J**). Based on these findings, we concluded that loss of Trim14 renders macrophages susceptible to apoptosis, consistent with previous links between Trim14 overexpression and hepatocellular and colorectal carcinomas ^38,40^.

Given that loss of Trim14 sensitizes macrophages to programmed cell death in response to Mtb infection and apoptotic triggers, we hypothesized that overexpression of Trim14 would reduce sensitivity to these triggers. To test this hypothesis, we generated doxycycline (Dox)-inducible RAW 264.7 macrophages overexpressing 3xFLAG-GFP or 3xFLAG-Trim14. We optimized the timing of doxycycline treatment, settling on 6h, to reduce off-target effects associated with prolonged doxycycline exposure (**Fig. S1I**). We found that 3xFLAG-Trim14 overexpression was sufficient to significantly dampen Mtb-induced and ABT-737 triggered cell death (**Fig. 1K-M**). DOX-inducible iBMDMs overexpressing 3xFLAG-GFP or 3xFLAG-Trim14 showed the same phenotype (**Fig. S1J-L**). As expected, caspase-3 cleavage was reduced in 3xFLAG-Trim14 overexpressing iBMDMs post-ABT-737 stimulation by immunoblot (**Fig. 1N**). Together, these data firmly implicate Trim14 in controlling the susceptibility of macrophages to undergo apoptotic cell death.

### Trim14 regulates mitochondria dynamics at rest and following apoptosis induction

We next wanted to understand the molecular mechanisms underpinning Trim14’s role in Mtb-and ABT-737-triggered apoptosis. Because alterations in the expression of pro- and anti-apoptotic genes can influence sensitivity to apoptotic stimuli and is frequently observed during infection and in cancer^51,52^, we first wanted to test if loss of Trim14 results in transcriptional reprogramming of macrophages. We performed RNA-seq on WT and *Trim14^-/-^* BMDMs at rest and at 4h post-Mtb infection and analyzed differences in relative transcript abundance by DESeq2. Overall, we observed few significant transcriptional changes in *Trim14^-/-^* cells in either condition (**Fig. 2A-B**) and RT-qPCR confirmed that loss of Trim14 does not dramatically impact expression of anti-apoptotic genes (*Bcl2* and *Bcl2l1*) or the pro-apoptotic gene *Bax* during Mtb infection (**Fig. S2A**) or after ABT-737 treatment (**Fig. S2B**). Protein levels of Bax and Bcl-xl were also unchanged between the two genotypes (**Fig. S2C**). Apoptotic gene expression at the transcript or protein level was similarly unchanged in cells overexpressing Trim14 (**Fig. S2D-F**). Thus, we concluded that gene expression differences were unlikely to be influencing susceptibility to apoptosis in *Trim14^-/-^* or Trim14 overexpression macrophages.

**Figure 2.**
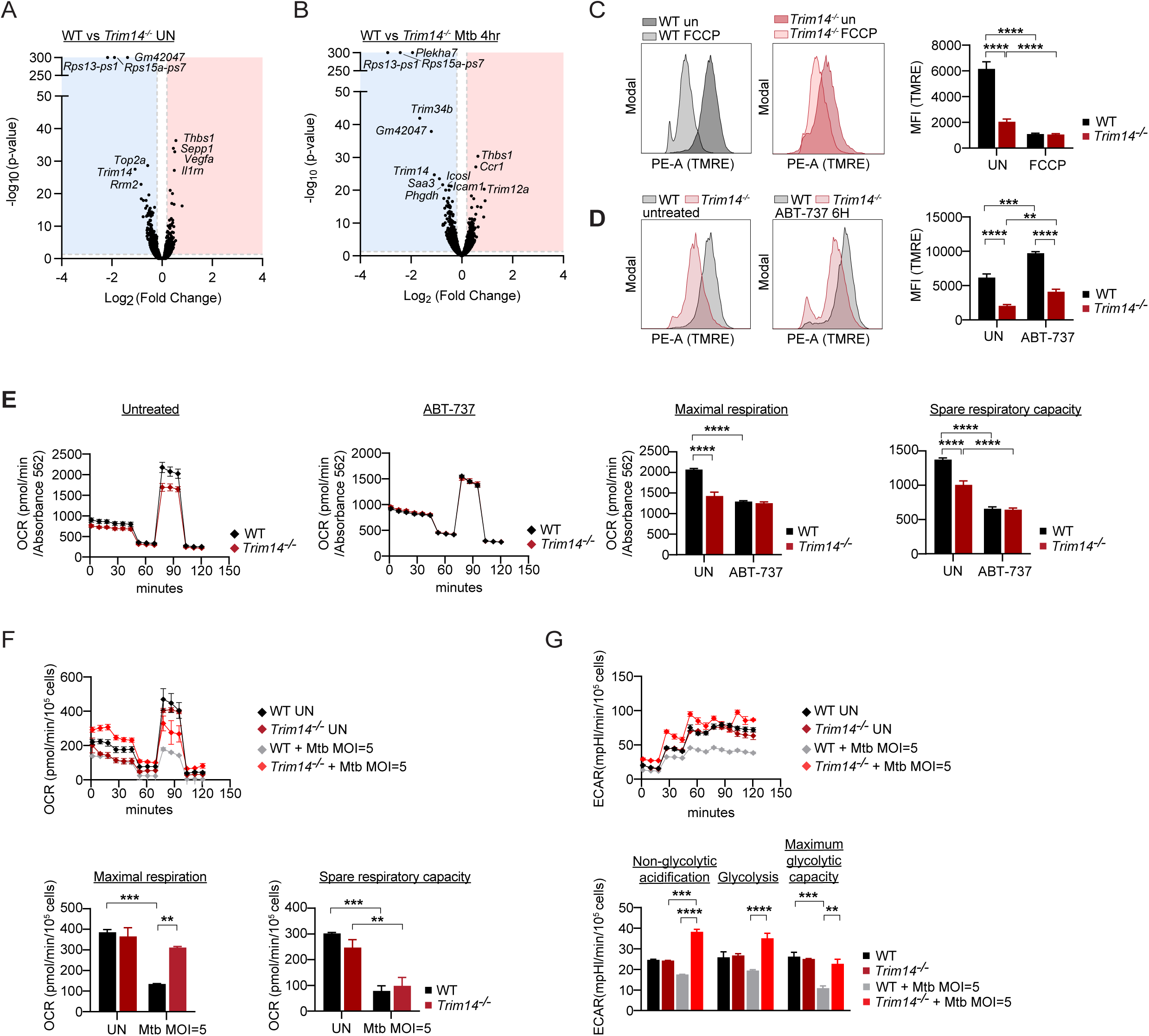
Trim14 is required to maintain mitochondrial homeostasis during Mtb infection. A) Volcano plot of genes differentially expressed in untreated WT and *Trim14^-/-^* BMDMs. Blue genes on the left are downregulated and red genes on the right are upregulated in *Trim14^-/-^*. B) As in (A) but in 4hr Mtb infected (MOI=5) WT and *Trim14^-/-^* BMDMs. C) Flow cytometry analysis of TMRE in FCCP treated (50 µM for 15 min) WT and *Trim14^-/-^* iBMDMs. D) As in (C) but analysis in untreated and 6hr treatment of 10 µM ABT-737 in WT and *Trim14^-/-^* iBMDMs. E) Oxygen consumption rate (OCR) measured by Agilent Seahorse Metabolic Analyzer in WT and *Trim14^-/-^* BMDMs. Untreated and ABT-737 treated (10 µM 6hr) on left. Maximal respiration and spare respiratory analysis on right. F) As in (E) but with untreated and Mtb-infected (24h MOI=5) on top. Maximal respiration and spare respiratory analysis on bottom. G) Extracellular acidification rates (ECAR) measured by Agilent Seahorse Metabolic Analyzer in WT and *Trim14^-/-^* BMDMs. Untreated and Mtb-infected (24h MOI=5) on top. Non-glycolytic acidification, glycolysis, and maximum glycolytic capacity analysis on bottom. Statistical analysis: *p < 0.05, **p < 0.01, ***p < 0.001, ****p < 0.0001. Statistical differences were determined for (C-G) using two-way ANOVA with Tukey’s post-test.

Because mitochondria serve as major signaling hubs to execute the final steps of apoptosis and Trim14 has been previously shown to localize to mitochondria^36^, we surmised that sensitivity to apoptosis in *Trim14^-/-^* macrophages could result from disruption of mitochondrial homeostasis. To measure the overall health of mitochondria in *Trim14^-/-^* and WT macrophages, we measured membrane potential using the cell-permeant dye tetramethylrhodamine (TMRE), which accumulates in healthy, uncompromised mitochondria. We found that *Trim14^-/-^* macrophages exhibit reduced membrane potential at rest (**Fig. 2C**) and following ABT-737 treatment (**Fig. 2D**). Despite previously described correlations between membrane potential and ROS production^53^, we did not observe any changes in mitochondrial ROS via mitoSOX in untreated or ABT-737 treated WT vs. *Trim14^-/-^* macrophages (**Fig. S2G**), nor did we detect differences in transcript abundance of the interferon stimulated gene *Ifit1* at rest or following ABT-737 treatment (**Fig. S2H**). These data argue that although mitochondrial membrane potential is compromised in *Trim14^-/-^* BMDMs, this defect is not sufficient to release mtDNA or activate cGAS-STING signaling^54,55^.

We next asked how loss of Trim14 affected mitochondrial bioenergetics using the Agilent Seahorse Metabolic Analyzer. In untreated cells, we measured a modest decrease in spare respiratory capacity and maximal respiration (**Fig. 2E)** but no change in glycolytic capacity in *Trim14^-/-^* macrophages compared to WT (**Fig. S2I**). While ABT-737 treatment decreased spare respiratory capacity overall, there were again, no significant differences between genotypes and no differences in ECAR (**Fig. 2E, S2J**). We then asked how *Trim14^-/-^* macrophages reprogram bioenergetics in response to Mtb infection. Typically, Mtb induces a decrease in maximal respiration, spare respiratory capacity, and glycolysis as infected macrophages enter an initial quiescent-like state^26^. This reprogramming, as measured by both OCR and ECAR, was almost completely absent in *Trim14^-/-^* BMDMs (**Fig. 2F-G**). However, the macrophage response to LPS, characterized by the Warburg shift from OXPHOS to glycolysis, remained intact in *Trim14^-/-^*macrophages (both after 4 and 16h treatment) (**Fig. S2K-L**), suggesting that there is something unique about mitochondria bioenergetic reprogramming in response to Mtb that requires Trim14.

### Trim14-dependent localization of Stat3 to mitochondria is required to maintain mitochondria homeostasis and control apoptosis

Having observed loss of mitochondrial membrane potential and failure to respond to the energy demands of Mtb infection, we wanted to know more about how Trim14 interacts with proteins involved in mitochondrial homeostasis. Our previous work found that Trim14 can physically interact with Tbk1 and Stat3, one of its substrates ^56^. We also found that loss of Trim14 results in higher levels of p754 Stat3, an inhibitory mark that can be deposited by Tbk1, following ISD transfection^37^ and Mtb infection (**Fig. S3A**). To ask whether Trim14 tunes apoptosis sensitivity through Stat3, we utilized Stattic, a small-molecule inhibitor of Stat3 activation and dimerization^57^. We found that Stattic increased apoptosis in WT BMDMs to *Trim14^-/-^* levels, but had no effect on *Trim14^-/-^* cell death (**Fig. 3A**). Because Stattic treatment phenocopies loss of Trim14 but has no effect in *Trim14^-/-^* cells, we surmised that enhanced apoptosis in *Trim14^-/-^*BMDMs was Stat3-dependent.

**Figure 3.**
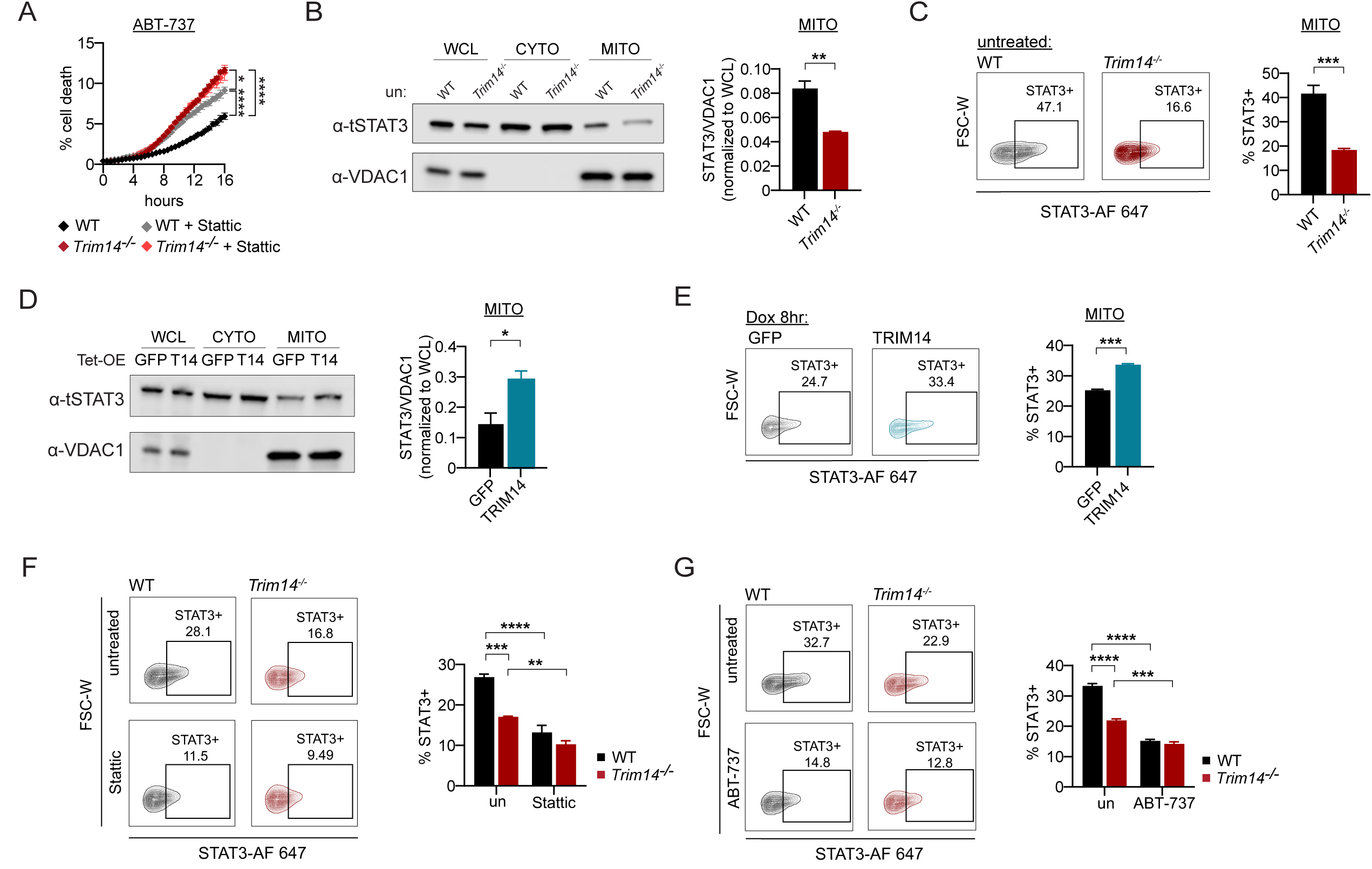
Trim14-dependent localization of Stat3 to mitochondria is required to maintain mitochondria homeostasis and control apoptosis. A) Cell death in WT and *Trim14^-/-^* BMDMs treated with 10 µM ABT-737 in combination with 5 µM Stattic. Cell death was measured by propidium iodide uptake (% cell death = propidium iodide positive/total cell count × 100%). B) Stat3 mitochondrial association measured by biochemical fractionation and immunoblot of untreated WT and *Trim14^-/-^* macrophages. Vdac1 was probed to control for mitochondria enrichment. Quantification of Stat3 relative to Vdac1 normalized to whole cell lysate on right; n = 3. C) Mitoflow analysis to measure Stat3 mitochondrial association in untreated WT and *Trim14^-/-^* macrophages. Quantification of %Stat3+ mitochondria shown on right. D) As in (B) but with tetracycline-inducible overexpression iBMDMs expressing 3X-Flag GFP or 3X-Flag TRIM14 for 8hr. E) As in (C) but with tetracycline-inducible overexpression iBMDMs expressing 3X-Flag GFP or 3X-Flag TRIM14 for 8hr F) Mitoflow analysis to quantify Stat3 mitochondrial association in WT and *Trim14^-/-^* macrophages treated with 10 µM Stattic for 6h. Quantification of %Stat3+ mitochondria on right. G) As in (F) but with WT and *Trim14^-/-^* macrophages treated with 10 µM ABT-737 for 6h. Statistical analysis: *p < 0.05, **p < 0.01, ***p < 0.001, ****p < 0.0001. Statistical differences were determined for (A, F, G) using two-way ANOVA with Tukey’s post-test and for (B, C, D, E) using unpaired two-tailed Student’s t test.

Our next question was how Stat3 function might be altered in *Trim14^-/-^* cells. Having measured few transcriptional differences in *Trim14^-/-^* BMDMs (**Fig. 2A-B, S2A-D**), we concluded that Stat3’s role in Trim14-regulated apoptosis was unlikely to stem from its role as a transcription factor. Previous studies have reported that a small pool of Stat3 localizes to mitochondria (mito-Stat3), where it is thought to regulate electron transport chain (ETC) function and mitochondrial permeability transition pore dynamics through cyclophilin D^58^. To test if Trim14 regulates Stat3 subcellular localization, we performed biochemical fractionation to enrich mitochondria from WT and *Trim14^-/-^* BMDMs. Immunoblot analysis revealed that mitochondrial Stat3 levels were significantly lower in *Trim14^-/-^* mitochondria than in WT BMDMs (**Fig. 3B**). To confirm these findings, we adopted a flow cytometry-based approach (mito-FLOW) using MitoView Green to label mitochondria independently of membrane potential, together with Stat3 antibody staining after permeabilization to quantify total protein levels^27,59,60^. To confirm assay specificity, we analyzed mitochondria isolated from *Stat3^-/-^* RAW 264.7 macrophages, which showed minimal off-target staining (**Fig S3B-D**). We found that mito-Stat3 levels were markedly reduced in untreated *Trim14^-/-^* macrophages when compared with WT controls (**Fig. 3C**). Conversely, 3xFLAG-Trim14 overexpression in iBMDMs increased mito-Stat3 levels by both immunoblot and mito-FLOW (**Fig. 3D-E**).

We next wanted to know if inhibiting Stat3 activity could impact its mitochondrial targeting. Stattic works by binding to the SH2 domains of Stat3, blocking dimerization and limiting phosphorylation at Y705 and S727. Treatment of WT macrophages with Stattic markedly decreased mito-Stat3 abundance (**Fig. 3F**), suggesting that dimerization and/or phosphorylation of Stat3 controls its mitochondrial localization and/or import. Interestingly, induction of apoptosis via ABT-737 treatment also decreased mito-Stat3 levels in WT macrophages, bringing them to levels comparable to those observed in *Trim14*-/- macrophages (**Fig. 3G)**. This finding suggests that Stat3 is relocalized from the mitochondria to the cytosol as cells initiate apoptosis, akin to releasing a brake. This decrease is not as dramatic in *Trim14^-/-^* macrophages, consistent with the brake already being released. Together, these data demonstrate that Trim14 controls the levels of Stat3 in the mitochondria and hint at a protective role for mito-Stat3 in limiting apoptosis following ABT-737 treatment.

To functionally link apoptosis sensitivity with Stat3 mitochondrial localization, we ectopically expressed either 3xFLAG-STAT3 or a mitochondria-targeted variant (3xFLAG-MTS-STAT3) in WT iBMDMs (**Fig. 4A**)^61^. In this construct, a MTS tag (the COX8A mitochondrial targeting pre-sequence (MSVLTPLLLRGLTGSARRLPVPRAK)) directs Stat3 to mitochondria, while DNA-binding domain mutations (VVV461–463AAA and EE434–435AA) prevent canonical Stat3 transcriptional activity, allowing us to ask if enforced mitochondrial targeting of Stat3 is sufficient to limit apoptosis^49,62,63^. To first test if this construct efficiently targets Stat3 inside mitochondria, we performed a proteinase K protection assay as in^64,65^. Unlike the outer mitochondrial membrane protein Tom20, whose surface-exposed domain is cleaved by proteinase K, a portion of 3xFLAG-STAT3 and 3xFLAG-MTS-STAT3, as well as the resident inner mitochondrial membrane protein NDUFB8 were proteinase K resistant (**Fig. 4B**), consistent with Stat3 accumulating inside mitochondria. We next asked how targeting Stat3 to the mitochondria impacts cell death. We found that while overexpression of 3xFLAG-STAT3 was sufficient to reduce cell death in iBMDMs, cell death was further reduced by 3xFLAG-MTS-STAT3 (**Fig. 4C**). These findings support a model whereby Trim14 directs Stat3 mitochondrial localization and the levels of mito-Stat3 set the threshold for macrophage apoptosis.

**Figure 4.**
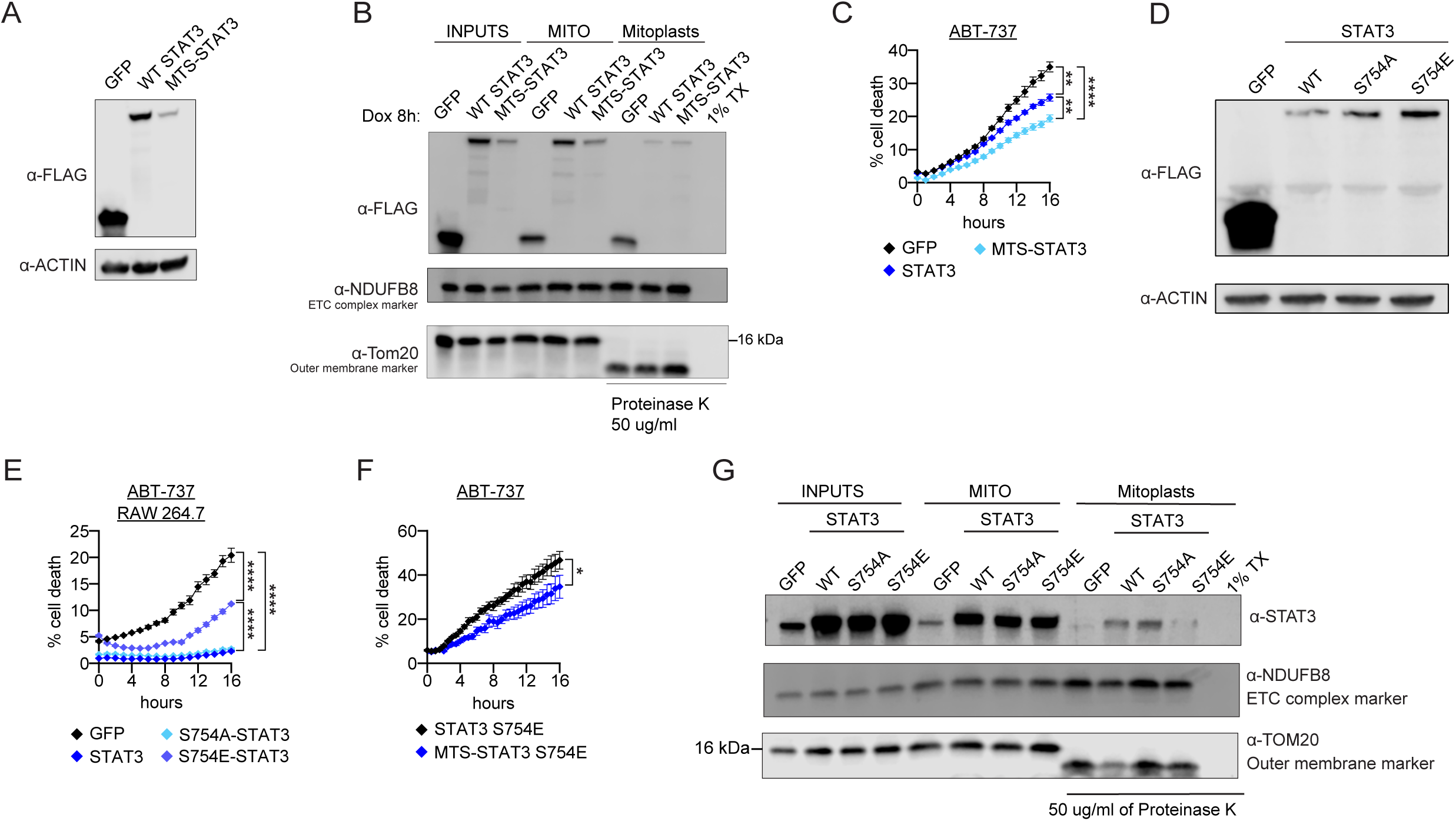
pS754 Stat3 limits Stat3 mitochondrial targeting and increases macrophage cell death. A) Immunoblot untreated iBMDMs stably expressing 3X-Flag GFP, 3X-Flag STAT3, or 3X-Flag MTS-STAT3. Actin was used as a loading control. B) Biochemical fractionation and immunoblot of tetracycline-inducible overexpression iBMDMs expressing 3X-Flag GFP, 3X-Flag STAT3, or 3X-Flag MTS-STAT3 for 8hr. Anti-Flag antibody was used to measure mitochondria association before and after treatment with 50µg/ml proteinase K (PK) to remove cytosolic contaminants from the mitochondria. Ndufb8 is an inner mitochondrial protein protected from PK. Outer membrane protein Tom20 is degraded upon PK treatment. C) Cell death in iBMDMs stably expressing 3X-Flag GFP, 3X-Flag STAT3, or 3X-Flag MTS-STAT3 treated with 10 µM ABT-737. D) Immunoblot analysis of untreated RAW 264.7 macrophages stably expressing 3X-Flag GFP, 3X-Flag STAT3, 3X-Flag STAT3 S754A, or 3X-Flag STAT3 S754E. Actin was used as a loading control. E) As in (C) but with RAW 264.7 macrophages expressing 3X-Flag GFP, 3X-Flag STAT3, 3X-Flag STAT3 S754A, or 3X-Flag STAT3 S754E. F) Cell death in tetracycline-inducible iBMDMs expressing 3X-Flag-STAT3 S754E and the corresponding MTS-tagged construct following doxycycline induction (1 µg/mL, 8h) and 5µM ABT-737. Actin was used as a loading control. G) Biochemical fractionation and immunoblot analysis of untreated iBMDMs stably expressing 3X-Flag GFP, 3X-Flag STAT3, 3X-Flag STAT3 S754A, or 3X-Flag STAT3 S754E. Stat3 was probed to measure mitochondria association before and after treatment with 50µg/ml proteinase K (PK) to remove cytosolic contaminants from the mitochondria. Ndufb8 and Tom20 serve as controls as in (B). Statistical analysis: *p < 0.05, **p < 0.01, ***p < 0.001, ****p < 0.0001. Statistical differences were determined for (C, E, F) using two-way ANOVA with Tukey’s post-test.

Because we have previously reported that Trim14 interacts with Stat3 and the kinase Tbk1^37^ and have repeatedly observed higher levels of pS754 Stat3 in *Trim14^-/-^* macrophages^37^ (**Fig. S3A**), we hypothesized that Trim14-dependent post-translational modification of Stat3 may direct its subcellular localization. To test the impact of the S754 residue on apoptosis and mitochondrial recruitment, we used site-directed mutagenesis to generate and overexpress a phospho-dead (S754A) and phospho-mimetic (S754E) variant of Stat3 (**Fig. 4D**) and stably expressed them in RAW 264.7 cells. While ectopic expression of 3xFLAG-STAT3 and 3xFLAG-STAT3 S754A conferred protection against ABT-737-induced apoptosis, the 3xFLAG-STAT3 S754E mutation limited this protective effect (**Fig. 4E**). Importantly, when we targeted the 3xFLAG-STAT3 S754E construct to mitochondria via an MTS, apoptosis was decreased relative to untargeted S754E (**Fig. 4F, S3F**), suggesting that Stat3-S754E cannot get to the mitochondria, but can still do its job if sent there. To confirm that phosphorylation at S754 alters mito-Stat3 abundance, we performed mito-FLOW analysis with WT, S754A, and S754E Stat3- expressing cells (**Fig. S3E**). Although total mitochondrial localization was comparable between the cells expressing the three Stat3 alleles (middle “MITO” lanes, **Fig. 4G**), proteinase K treatment significantly reduced levels of 3xFLAG-STAT3 S754E, indicating diminished import or retention within mitochondria (right “Mitoplasts” lanes, **Fig. 4G**). Together, these data suggest that phosphorylation of Stat3 at S754 reduces its mitochondrial localization and limits its ability to protect against apoptosis.

### Mitochondrial-associated Stat3 controls apoptosis via mPTP dynamics

We next sought to define the mechanism through which Stat3 mitochondrial recruitment alters apoptosis sensitivity in macrophages. Mito-Stat3 has previously been shown to interact with Cyclophilin D (CypD), a mitochondrial peptidyl-prolyl cis-trans isomerase that acts as a key regulator of the mitochondrial permeability transition pore (mPTP). To test if Stat3’s ability to protect against apoptosis occurs through regulation of the mPTP, we used an inhibitor of CypD, cyclosporin A (CsA), to inhibit its activity and induce mPTP closure^66,67^. We found that CsA reduced cell death following ABT-737 treatment in WT cells as expected and returned the enhanced cell death in *Trim14^-/-^* BMDMs back to WT levels (**Fig. 5A**). Having implicated the mPTP in Trim14-dependent cell death, we next employed a Calcein AM-Cobalt (II) chloride (CoCl_2_) assay to directly measure mPTP dynamics. Briefly, WT vs *Trim14^-/-^* iBMDMs were loaded with the non-fluorescent dye Calcein AM, which is fluorescent in live, intact cells^68,69^. When cells are healthy and the mPTP is closed, CoCl_2_ quenches calcein in the cytosol but cannot reach the mitochondria. If the mPTP is open, CoCl_2_ can access the mitochondria and quench calcein signal there as well (**Fig. S4A**). We found that *Trim14^-/-^* iBMDMs had decreased calcein signal after quenching, suggesting mPTPs are more open in *Trim14^-/-^* mitochondria (**Fig. 5B**). This trend was maintained during Mtb infection of WT and *Trim14^-/-^* iBMDMs (**Fig. 5C**). Consistent with our finding that ABT-737 treatment promotes what looks like release of mito-Stat3, ABT-737 treatment enhanced mPTP opening in both WT and *Trim14^-/-^* iBMDMs whereas CsA closed pores, maintaining calcein signal (**Fig. 5D**). Overexpression of Trim14 limited mPTP opening (**Fig. 5E, S4B**), consistent with earlier observations that Trim14 overexpression limits cell death in macrophages (**Fig. 1K-M**). Also consistent with earlier cell death measurements (**Fig. 4C**), overexpression of 3X-FLAG STAT3 and 3X-FLAG-MTS-STAT3 dramatically decreased mPTP opening (**Fig. 5F, S4C**). Collectively, these data suggest that the Trim14-Stat3 axis limits apoptosis by controlling mPTP dynamics.

**Figure 5.**
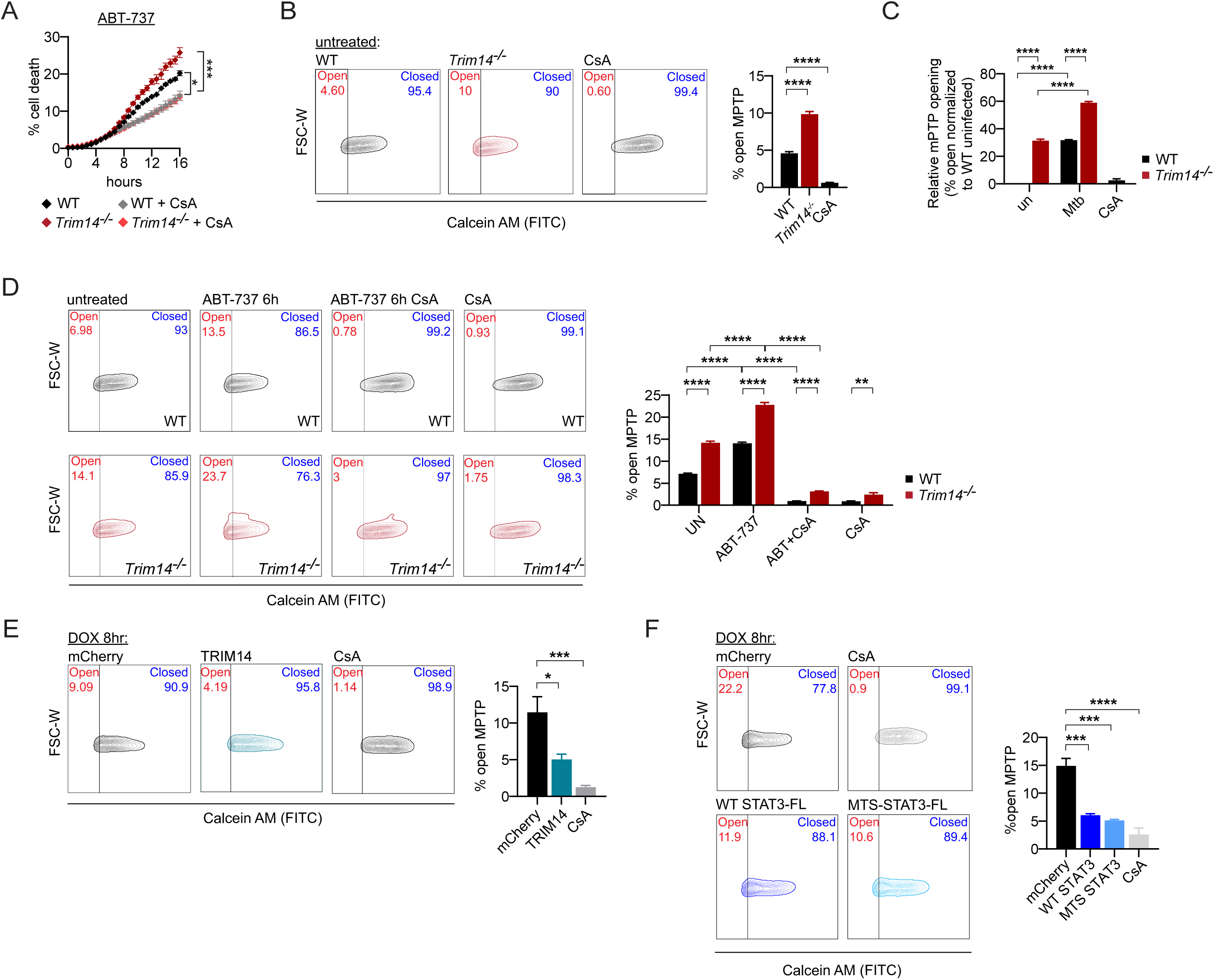
Trim14 reduces mPTP opening by maintaining mito-Stat3 levels. A) Cell death in WT and *Trim14^-/-^* macrophages treated with 10 µM ABT-737 and 5 µM CsA. Cell death was measured by propidium iodide uptake (% cell death = propidium iodide positive/total cell count × 100%). B) Flow cytometry analysis of calcein fluorescence after CoCl_2_ quench in untreated WT and *Trim14^-/-^* macrophages. WT macrophages were treated for 16h with 0.5 µM CsA to reduce mPTP opening. Quantification shown on right. C) Calcein fluorescence measured by microplate reader after CoCl_2_ quench in uninfected and Mtb-infected (MOI=5) WT and *Trim14^-/-^* macrophages for 6h. WT macrophages were treated for 16h with 0.5 µM CsA to reduce mPTP opening. Total cell count was measured using propidium iodide in 0.1% Triton-X (100% cell death). PI-normalized calcein fluorescence was expressed relative to WT uninfected cells, relative mPTP opening was calculated as percent loss of normalized calcein signal. D) As in (B), but with untreated, 10 µM ABT-737, 10 µM ABT-737 + 5 µM CsA, and 0.5 µM CsA (16h) at 6h post-apoptotic induction in WT and *Trim14^-/-^* macrophages. Quantification shown on right. E) As in (B), but with tetracycline-inducible iBMDMs expressing mCherry or 3X-Flag TRIM14 for 8hr. mCherry over-expressing iBMDMs were treated for 16h with 0.5 µM CsA to reduce mPTP opening. Quantification shown on right. F) As in (E), but with tetracycline-inducible iBMDMs expressing mCherry, 3X-Flag STAT3, or 3X-Flag MTS-STAT3 for 8hr. mCherry over-expressing iBMDMs were treated for 16h with 0.5 µM CsA to reduce mPTP opening. Quantification shown on right. Statistical analysis: *p < 0.05, **p < 0.01, ***p < 0.001, ****p < 0.0001. Statistical differences were determined for (B, E, F) using one-way ANOVA with Tukey’s post-test and (A, C, D) using two-way ANOVA with Tukey’s post-test.

### Loss of Trim14 enhances host resistance to *in vivo* Mtb infection

Our *in vitro* macrophage experiments demonstrate that Trim14 negatively regulates macrophage apoptosis by targeting Stat3 to mitochondria and controlling opening of the mPTP. Given that cell death modality usage is a decisive factor in tuberculosis pathogenesis—whereby apoptosis promotes bacterial clearance over pro-inflammatory necrosis^3^—we hypothesized that increased apoptosis due to Trim14 deficiency would enhance host resistance to Mtb *in vivo*.

To first assess the ability of WT and *Trim14*^⁻/⁻^ mice to survive Mtb infection, we infected male and female mice from both genotypes with ∼750 Erdman bacilli by aerosol delivery. We deliberately selected a medium-dose inoculum for this experiment to speed disease progression and impose a stringent challenge on the *Trim14*^⁻/⁻^ mice. We found that *Trim14^-/-^* mice are more resistant to Mtb infection, with a median survival time of 123 days compared to 84 days in WT mice (**Fig. 6A**). Increased survival correlated with decreased weight loss (**Fig. 6B**). Consistent with female mice being more resistant to Mtb in general^70^, we measured higher median survival times in females (144 days) compared to males (109 days) (**Fig. S5A-C**).

**Figure 6.**
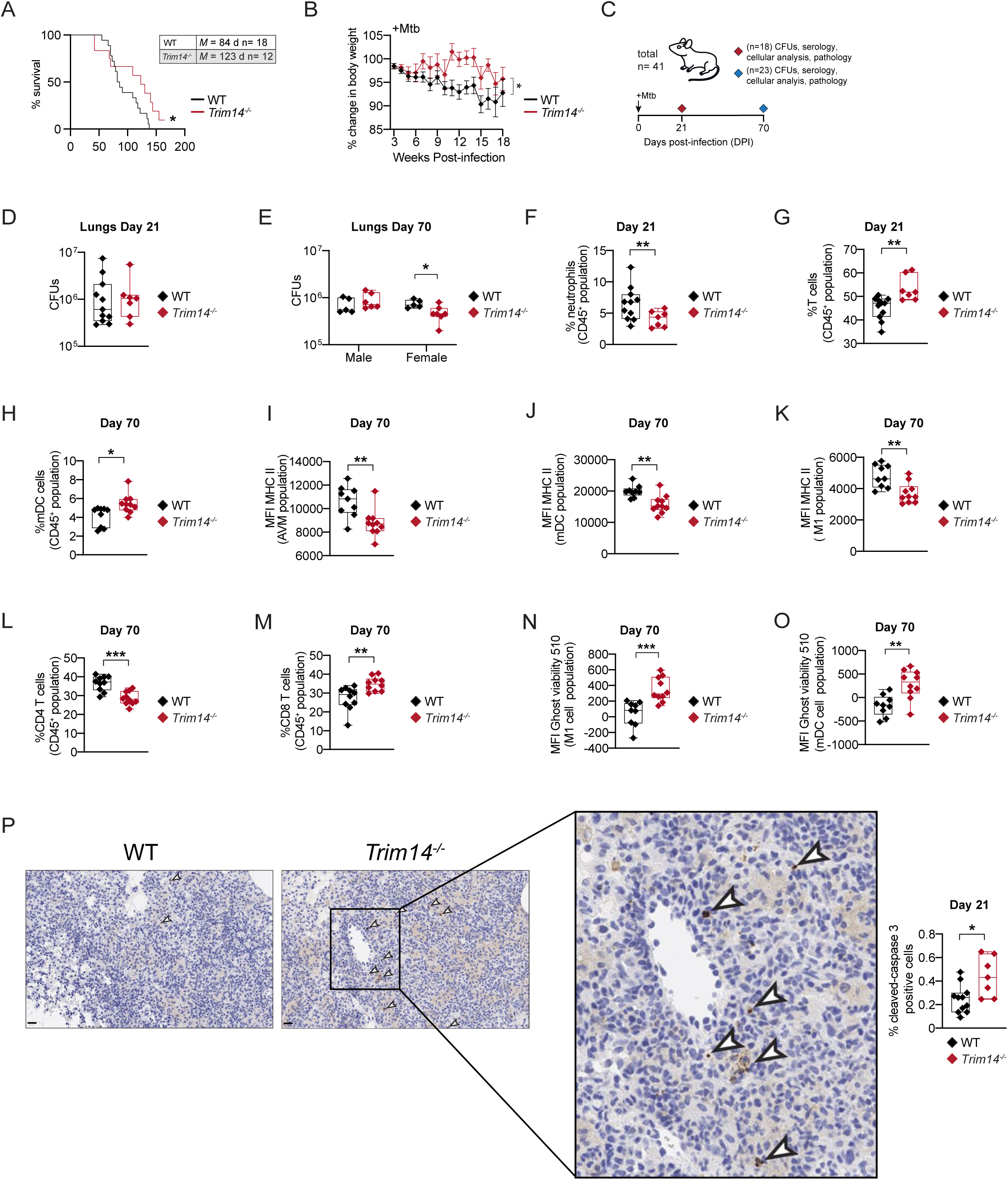
Loss of Trim14 enhances host resistance to Mtb infection *in vivo*. A) Survival analysis of WT (n=18) and *Trim14^-/-^* (n=12) mice infected with ∼750 Mtb bacilli. B) Percent body weight change of WT (n=18) and *Trim14^-/-^* (n=12) mice infected with ∼750 Mtb bacilli. C) Experimental timeline of WT and *Trim14^-/-^* mice infected with a ∼100 Mtb bacilli at day 21 and day 70. D) Lung CFUs of WT and *Trim14^-/-^* mice at day 21. E) Lung CFUs of WT and *Trim14^-/-^* mice at day 70 separated by sex. F) FACs analysis of neutrophils expressed as a % of total CD45⁺ in WT and *Trim14^-/-^* mice lungs at day 21. G) As in (F) but measuring the amount of T cells expressed as a % of total CD45⁺. H) FACs analysis of dendritic cells expressed as a % of total CD45⁺ in WT and *Trim14^-/-^* mouse lungs at day 70. I) Mean fluorescence intensity (MFI) of MHC class II expression in alveolar macrophages from WT and *Trim14^-/-^* mouse lungs at day 70. J) As in (I) but MFI of dendritic cells. K) As in (I) but MFI of M1 macrophages. L) FACs analysis of CD4 T cells expressed as a % of total CD45⁺ in WT and *Trim14^-/-^* mouse lungs at day 70. M) As in (L) but CD8 T cells. N) MFI of Ghost Viability 510 dye in M1 macrophages WT and *Trim14^-/-^* mouse lungs at day 70. O) As in (N) but MFI of dendritic cells. P) Representative images of cleaved-caspase 3 IHC in WT and *Trim14^-/-^* mouse lungs at day 21. Percent of positive cleaved-caspase 3 normalized to total cell count. Scale bar = 20 µM. Statistical analysis: *p < 0.05, **p < 0.01, ***p < 0.001, ****p < 0.0001. Statistical differences were determined for (A) Mantel-Cox log-rank, (B) two-way ANOVA with Tukey’s post-test (D-P) using Mann-Whitney U test. one-way ANOVA with Tukey’s post-test.

Having confirmed that loss of Trim14 confers protection against Mtb in a mouse model, we wanted to more closely examine the immune responses driving this protection. Briefly, we carried out a low-dose aerosol infection (∼100 bacilli) and sacrificed WT and *Trim14^-/-^* mice at day 21 and 70, to capture the culmination of the innate immune response and a fully engaged adaptive immune response, respectively (**Fig. 6C**). Overall, we did not see major differences in bacterial burdens in the lungs and spleens of *Trim14^-/-^* mice on day 21 (**Fig 6D, S5D**). There were, however, significantly fewer bacilli in the lungs of *Trim14^-/-^* female mice at day 70, consistent with enhanced protection in *Trim14^-/-^* females (**Fig. 6E, S5E**). Gross examination of lungs with H&E staining did not reveal major qualitative or quantitative differences in hallmarks of excessive pulmonary inflammation, including neutrophilic aggregates, necrotic debris, or necrotic inflammation, in *Trim14^-/-^* mice compared with WT controls. (**Fig. S5F-I**).

We next used flow cytometry to evaluate changes in lung immune cell populations in WT and *Trim14^-/-^* mice at day 21 and 70 (Gating strategy provided in **Fig. S6A-B**). At day 21, we measured a slight increase in the total number of CD45+ cells in *Trim14^-/-^* mice (**Fig. S5J**) and a slight decrease in % neutrophils (**Fig. 6F**), but no significant differences in alveolar macrophages, monocytes, monocyte-derived macrophages, or B cells (**Fig. S5K**). We did however, note a 20% increase in the %T cells (**Fig. 6G**), which play an obvious protective role in Mtb infection^6,7,10^. By day 70, CD45+ cells were the same in the two genotypes and no differences were detected in alveolar macrophage, neutrophil, monocyte, macrophage, or NK cell numbers (**Fig. S5L-O**). Consistent with growing evidence for B cells in protective antimycobacterial immunity^71^, we measured increased plasma cell numbers in *Trim14^-/-^* mice relative to WT (**Fig. S5P**). There was also a slight increase in dendritic cells (DCs) in *Trim14^-/-^*mice at day 70 (**Fig. 6H**). More striking, however, was a significant decrease in MHCII surface expression in multiple types of antigen presenting cells (alveolar macrophages, dendritic cells, and M1 macrophages) in *Trim14^-/-^* mice (**Fig. 6I-K**). Defects in MHCII levels prompted us to look more carefully at T cell populations at day 70. Consistent with lower MHCII expression, we observed fewer CD4^+^ T cells in *Trim14^-/-^* mice (**Fig. 6L**). This decrease in CD4+ T cells was concomitant with a sizeable increase in CD8^+^ T cells in *Trim14^-/-^* mice compared to WT controls (**Fig. 6M**). *Trim14^-/-^* mice also had increased number of memory T cells (both CD4^+^ and CD8^+^) (**Fig. S5Q**). Overall, the increase in DCs, decrease in MHCII surface expression on APCs, decrease in CD4^+^ T cells, and increase in CD8^+^ T cells, are consistent with a model whereby DC-mediated cross-presentation of ingested apoptotic bodies enhances protective CD8-mediated antimycobacterial immunity in *Trim14^-/-^* mice.

This model hinges on there being more apoptosis in the lungs of *Trim14^-/-^* mice. To measure this directly, we used an amine-reactive fluorescent Ghost viability dye to quantify cell death in different immune cell populations by flow cytometry. Consistent with our *in vitro* findings (**Figs. 1-5**), we found that M1 macrophages and dendritic cells had decreased viability in *Trim14^-/-^* mice (**Fig 6N-O)**. Consistent with decreased innate immune cell viability, we observed a significant increase in apoptosis in *Trim14^-/-^* mouse lungs as measured by cleaved-caspase 3 via immunohistochemistry (**Fig. 6P**). Together, this work identifies Trim14 as an important negative regulator of antimycobacterial immunity and suggests that enhancing the specific type of apoptosis favored in *Trim14^-/-^* cells may help boost antibacterial immunity *in vivo*.

These *in vivo* findings raised the possibility that increased apoptosis of *Trim14^-/-^* cells may enhance antigen transfer via cross presentation and subsequent CD8^+^ T cell activation. Importantly, this effect was unlikely to reflect pre-existing differences in the peripheral T cell compartments, as naïve WT and *Trim14^-/-^* mice exhibited comparable numbers of CD4^+^ and CD8^+^ T cells in the spleen and lymph nodes (**Fig. S7A-B**). To model this process *ex vivo*, we infected WT macrophages with Mtb-OVA, a strain expressing OVA peptide sequences fused to the secreted Mtb virulence factor CFP-10, which enables OVA-derived antigen presentation through MHC class I and II pathways (**Fig. 7A-B, S7C**). Infected macrophages were then co- cultured with OT-I splenocytes, whose CD8^+^ T cells recognize the OVA_257–264_ peptide presented on MHC class I. We validated this system by comparing WT Mtb Erdman, Mtb-OVA Erdman, and SIINFEKL peptide-treated macrophage co-cultures. Mtb-OVA-infected macrophages increased CD69 expression on OT-I CD8^+^ T cells compared with WT Mtb-infected controls, confirming antigen-specific CD8^+^ T cell activation in this *ex vivo* system (**Fig. 7B, S7D**). We then asked whether Trim14 loss alters the capacity of infected macrophages to promote antigen-specific CD8^+^ T cell activation. To test this, WT and *Trim14^-/-^* macrophages were infected with Mtb-OVA and co-cultured with OT-I splenocytes, after which CD69 expression was measured on CD8^+^ T cells. OT-I CD8^+^ T cells co-cultured with Mtb-OVA-infected *Trim14^-/-^* macrophages exhibited increased CD69 expression compared with those co-cultured with infected WT macrophages, suggesting that Trim14 deficiency enhances antigen-dependent CD8^+^ T cell activation (**Fig. 7C, S7F**).

**Figure 7.**
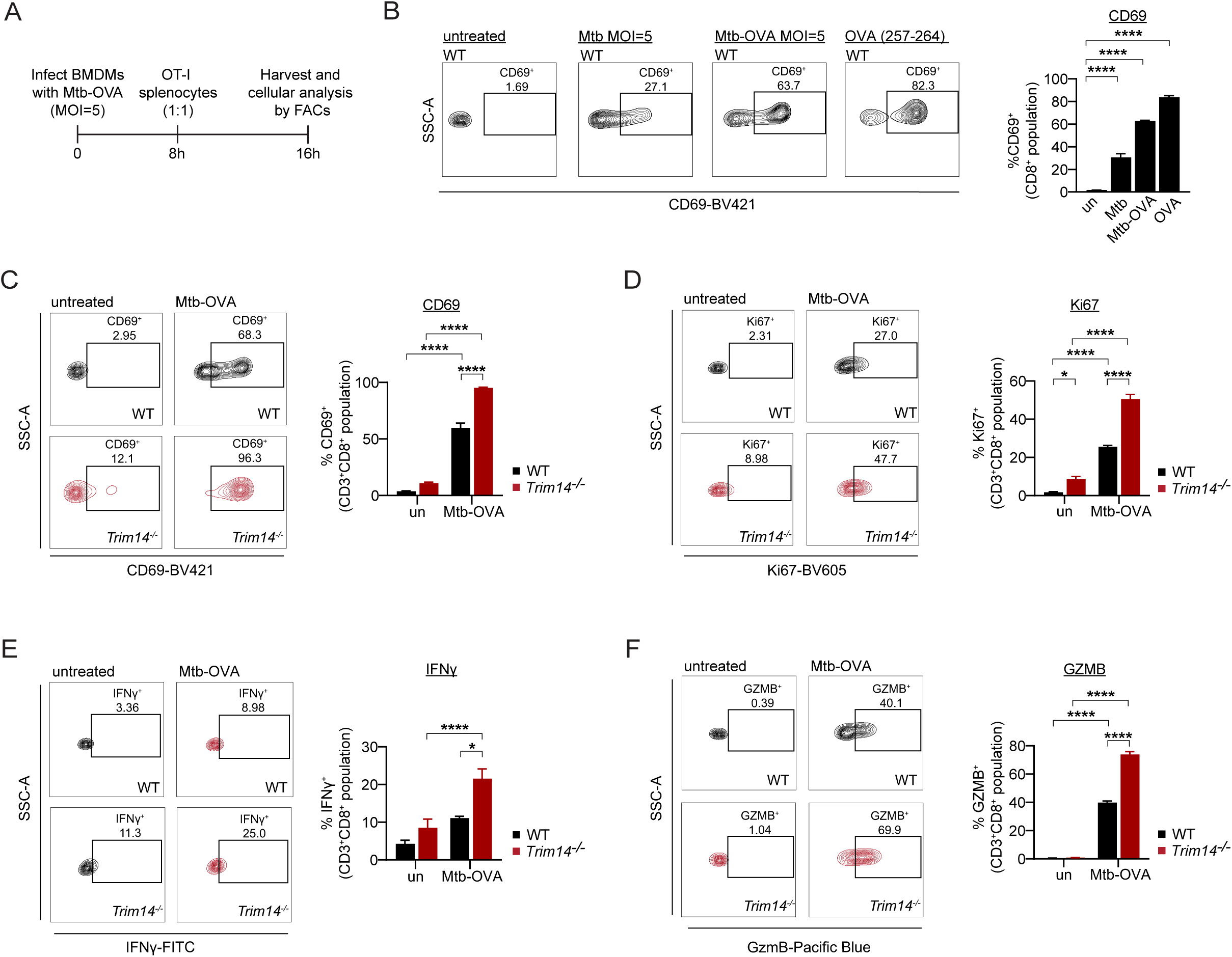
Trim14-deficient macrophages potentiate CD8^+^ T cell activation during Mtb infection. A) Experimental schematic of macrophages infected with Mtb-OVA and co-cultured with OT-I splenocytes B) FACs analysis of %CD69^+^ surface staining on CD8^+^ OT-I cells co-cultured with uninfected, WT Mtb (MOI=5), Mtb-OVA (MOI=5), and OVA peptide (257-264) stimulated iBMDMs. C) FACs analysis of %CD69^+^ surface staining on CD3^+^ CD8^+^ OT-I cells co-cultured with uninfected or Mtb-OVA (MOI=5) infected WT and *Trim14^-/-^* iBMDMs. D) As in (C) but measuring % Ki67^+^ CD3^+^ CD8^+^ OT-I cells. E) As in (C) but measuring % IFNγ^+^ CD3^+^ CD8^+^ OT-I cells. F) As in (C) but measuring % GZMB^+^ CD3^+^ CD8^+^ OT-I cells. Statistical analysis: *p < 0.05, **p < 0.01, ***p < 0.001, ****p < 0.0001. Statistical differences were determined for (B-F) using one-way ANOVA with Tukey’s post-test.

To further define the functional state of these OT-I CD8^+^ T cells, we performed intracellular staining for T cell proliferation and effector-associated markers. CD8^+^ T cells co-cultured with Mtb-OVA-infected *Trim14^-/-^* macrophages showed increased Ki67 expression, consistent with enhanced proliferative activity (**Fig. 7D, S7G**). Furthermore, these cells produced higher levels of IFNγ and granzyme B, suggesting that Trim14 loss promotes the development of antigen-specific CD8^+^ T cells with enhanced inflammatory and cytotoxic potential (**Fig. 7E-F, S7H-I**). Together, these findings suggest that the increased apoptotic susceptibility of *Trim14^-/-^* macrophages may enhance antigen availability for CD8^+^ T cell activation, thereby linking Trim14-dependent control of macrophage cell death to downstream adaptive immune responses during Mtb infection.

## DISCUSSION

Mitochondrial integrity represents a critical determinant of innate immunity, integrating metabolic homeostasis, inflammatory signaling, and cell death programs to shape host outcomes during *Mycobacterium tuberculosis* infection. This report identifies Trim14-dependent localization of mitochondrial Stat3 as an anti-apoptotic checkpoint that preserves mitochondrial integrity and limits apoptosis during Mtb infection. *In vivo*, loss of Trim14 promotes apoptosis of macrophages and dendritic cells, which amplifies CD8⁺ T cell responses and confers an approximately 30% improvement in survival compared to wild-type mice. These findings suggest that therapeutic disruption of the Trim14–Stat3 axis may enhance host-protective immunity against Mtb.

Collectively, our studies hint at two non-mutually exclusive mechanisms through which Trim14 regulates Stat3 mitochondrial targeting and function. The first possibility is that Trim14 primarily acts as a scaffold, physically bringing Stat3 to the mitochondria. We know that Trim14 and Stat3 can physically interact^37^, but where this complex forms in macrophages remains unexplored. Earlier studies of antiviral responses identified TRIM14 as a mitochondrial adaptor that localizes to the outer mitochondrial membrane and promotes assembly of the MAVS–NEMO/IKKγ signalosome^36^. It has also been shown to form a mitochondrial complex with WHIP and PPP6C that is required for efficient RIG-I-dependent signaling^36,72^. Since Trim14 can assemble protein complexes at mitochondria, it is possible it does the same for Stat3, bringing it nearby the outer membrane protein Tom20 and the ETC-associated protein GRIM19 (aka NDUFA13), which have been linked to Stat3 mitochondrial import^50,64^. Studies that make use of Trim14 mutants that fail to localize to the mitochondria (Trim14 R30A, L34A, and L46A)^36^ could help test this Trim14 handoff-import model.

The second possibility is that Trim14 control of Stat3 is driven not by its mitochondrial localization but by its ability to bind Tbk1 and direct the Stat3 residues it phosphorylates. Other studies have linked Tbk1 to phosphorylation of Stat3 at both S727 and S754^56,73^. Prior studies have linked Stat3 phosphorylation at S727 to mitochondrial Stat3 function and, in some contexts, to GRIM19-dependent mitochondrial import^61,64^. However, more recent work in S727A KI mice suggest that phosphorylation at S727 is dispensable for mitochondrial abundance and instead regulates Stat3 mitochondrial function once localized^74^. Our findings fit nicely with this newer model, whereby (1) Tbk1 preferentially phosphorylates Stat3 at S754 when Trim14 is absent, (2) pS754 negatively regulates mitochondrial targeting of Stat3, and (3) pS727 is important for controlling Stat3 activity once imported (**Fig. 4E-G**).

Our findings also raise questions about how mPTP opening is functionally coupled to specific cell death modalities during Mtb infection. In many contexts, sustained mPTP opening causes mitochondrial depolarization, bioenergetic collapse, and necrotic cell death^75,76^. For example, Mtb infection has been reported to promote pathogenic necrosis in macrophages in a TNF-driven mitochondrial ROS and calcium overload dependent manner. In contrast, our data suggest that mPTP opening in *Trim14^-/-^* macrophages preferentially lowers the threshold for apoptosis rather than driving necrosis. Importantly, despite altered mitochondrial function, *Trim14^-/-^* macrophages did not undergo the major bioenergetic collapse during Mtb infection that would be expected to favor necrotic cell death (**Fig. 2G-H**). These findings suggest that the consequences of mPTP opening during Mtb infection are context-dependent and may be shaped by the magnitude, timing, and duration of pore opening. This context dependence may be especially important when considering CypD as a therapeutic target during tuberculosis. Pharmacologic inhibition or genetic loss of CypD has been reported to limit mitochondrial permeability transition and necrotic cell death in Mtb-infected macrophages^77^. However, CypD-deficient mice are more susceptible to Mtb infection despite enhanced Mtb-specific T cell responses, suggesting that systemic loss of CypD can impair host protection through effects beyond macrophage necrosis, including altered T cell metabolism^78^. Together with our findings, these studies argue that CypD-dependent mPTP activity cannot be viewed as uniformly detrimental during tuberculosis but instead may have cell-type specific effects that impact whether host immunity is altered during Mtb infection.

A growing body of literature suggests that canonical STAT3 signaling favors Mtb persistence. In Mtb-infected human macrophages, STAT3 represses inflammatory cytokine production and nitric oxide synthesis and pharmacological inhibition of STAT3 limits Mtb replication^79^. In mice, loss of Stat3 in the myeloid compartment reduces bacterial burden, increasing IL-6 and IL-23 production and enhancing IL-17 responses from mycobacteria-specific CD4^+^ T cells^80^. Our report, which demonstrates that Stat3 constrains macrophage apoptosis, highlights yet another way that Stat3 represses antimycobacterial immunity. Together, these findings make a powerful argument for inhibition of Stat3 as an adjunct therapy to help control tuberculosis. Promoting apoptosis itself may also prove a useful antimycobacterial strategy, as combined BCL-2/MCL-1 inhibition and treatment with the BCL-2 inhibitor navitoclax^81,82^ was recently shown to reduce Mtb intracellular growth in human and murine macrophages and limit Mtb pathogenesis in mice.

Lastly, by revealing a function for Trim14 in regulating cell death programs, our findings extend the role of Trim14 beyond canonical innate immune signaling. This may be particularly relevant in cancer. Notably, TRIM14 expression has been repeatedly associated with breast cancer, colorectal cancer, and melanoma^38,39,83^, with several reports implicating STAT3 signaling in tumor progression. In this context, Trim14-dependent control of mitochondrial Stat3 may represent an additional mechanism through which cancer cells raise the threshold for apoptosis and acquire more aggressive or therapy-resistant phenotypes. Defining this axis across various disease settings may reveal new opportunities to modulate mitochondrial cell death programs for therapeutic benefit.

### Limitations of the study

Although our study identifies Trim14 as a regulator of mitochondrial Stat3 abundance, the precise mechanism by which Trim14 dictates Stat3 import or retention within mitochondria remains unresolved. Likewise, while we identify a mechanism in which Trim14 negatively regulates mPTP opening during apoptosis in a Stat3-dependent manner, we have not fully defined the mitochondrial binding partners through which Stat3 modulates pore dynamics. Intriguingly, a recent study found that immunoprecipitation of mitochondrial Stat3 enriched several proposed components of the mPTP, including adenine nucleotide translocases and ATP synthase subunits^84^. Future studies will be needed to determine whether Stat3 directly interacts with these proteins in Mtb-infected macrophages and whether these interactions are regulated by Trim14. Lastly, while *Trim14^-/-^* mice exhibit multiple hallmarks of host resistance to Mtb infection, the requirement for CD8+ T cells in this protective phenotype remains unresolved. CD8^+^ T cell depletion, adoptive transfer, or bone marrow chimera approaches will help define whether enhanced CD8^+^ T cell immunity is necessary and sufficient for protection in *Trim14^-/-^*mice.

## ACKNOWLEDGEMENTS

We would like to thank the members of the Watson and Patrick labs for providing invaluable feedback in review of this manuscript. We are grateful to Dr. JoAnne Flynn and Amy Fraser at University of Pittsburgh School of Medicine for providing the Mtb-OVA strain. Thanks to Dr. Cerdic Geoffroy (Texas A&M) for providing the original STAT3 constructs (MTS-tagged). We appreciate Dr. Joshua Bryant and the West lab (Texas A&M) for the Cre-J2 virus. We are especially grateful for the invaluable advice from Dr. Mary Philip (VUMC) on T cell co-culture assays. We are grateful for the technical expertise of Malea Murphy at the Integrated Microscopy and Imaging Laboratory and Robbie Moore at the Flow Cytometry & Cell Sorting Facility core facility at the Texas A&M Naresh K. Vashisht College of Medicine. The VMC Flow Cytometry Shared Resource is supported by the Vanderbilt Ingram Cancer Center (P30 CA68485) and the Vanderbilt Digestive Disease Research Center (DK058404). Translational Pathology Shared Resource is supported by NCI/NIH Cancer Center Support Grant P30CA068485. This work was supported by the NIH NIAID grants 5R01AI155621 (to ROW and KLP) and NIH F31 training grant 1F31AI176795 (CJM), 1F31AI176652 (AKC), 1F31CA298557 (MHS), and T32AI112541 (TMN).

## AUTHOR CONTRIBUTIONS

Conceptualization, R.O.W., C.J.M., K.L.P.; Investigation, C.J.M., A.K.C., M.H.S., S.L.H., T.M.N, M.J.C., L.W.S., C.G.W, K.L.P., and R.O.W.; Methodology, C.J.M., M.H.S., S.L.H., T.M.N, L.W.S., C.G.W, K.L.P., and R.O.W.; Writing, C.J.M., K.L.P., R.O.W.; Visualization, C.J.M., K.L.P., R.O.W.; Funding acquisition, C.J.M., A.K.C., M.H.S., T.M.N, K.L.P., and R.O.W.; Supervision, R.O.W. and K.L.P.

## Experimental Models and Subject Details

### Mouse husbandry and strains

*Trim14^-/-^* mice were developed by MMRRC at UC Davis using CRISPR Cas9 gene editing to constitutively delete exon 3-4 and flanking splicing regions in mouse zygotes (MMRC stock #043885-UCD). Subsequent founders were backcrossed to C57BL6/N (Jackson Laboratory strain #005304) to produce heterozygous animals. Genotyping was conducted using the following PCR primers: Trim14-comF (CAGGCTGATCTTGAACTTCCTGTGG), Trim14-wtR (GCTGTTCAAAGCAAGAGGCCACT, and Trim14-mutR (TCCTGGGCCGTAAGTTGTCAGCTA). WT PCR bands originated at 525 bp while mutant PCR bands at 394 bp (**Fig. S1A**). Loss of Trim14 protein was confirmed by western blot of BMDMs (**Fig. S1B**). C57BL/6-Tg (TcraTcrb)1100Mjb/J (OT-I) mice were purchased from Jackson Laboratory strain #003831. All mice used in experiments were compared to age- and sex- matched controls. Littermate controls were used in all experiments and fed a standard 4% chow diet. For *ex vivo* BMDM experiments, male mice 8-12 weeks old were used. For *ex vivo* T cell experiments, mice 8-12 weeks old were used. For *in vivo* experiments, mice were used at 10-12 weeks. All animals were housed, bred, and studied at Texas A&M Health Science Center and Vanderbilt University Medical Center under approved Institutional Animal Care and Use Committee guidelines.

### M. tuberculosis

The Erdman strain (Erdman WT, Erdman mCherry, Erdman OVA) was used for *M. tuberculosis* infections. Low passage lab stocks were thawed for each experiment to ensure virulence was preserved. *M. tuberculosis* was cultured in roller bottles at 37°C in Middlebrook 7H9 broth (BD Biosciences) supplemented with 10% OADC (BD Biosciences), 0.5% glycerol (Fisher), and 0.1% Tween-80 (Sigma) or on 7H10 plates. All work with *M. tuberculosis* was performed under Biosafety level 3 containment using procedures approved by the Texas A&M University and Vanderbilt University Medical Center Institutional Biosafety Committees.

### Primary and immortalized cell culture

Bone marrow derived macrophages (BMDMs) were differentiated from bone marrow isolated from mouse femurs and tibias with DMEM (HyClone) supplemented with 1mM sodium pyruvate. Cells were centrifuged at 400 rcf for 5 minutes and resuspended in BMDM media (DMEM with 1 mM sodium pyruvate, 20% heat inactivated FBS (Gibco), and 10% MCSF media (Watson lab)). Isolated BM cells were plated in 15 cm non-TC dishes at 5.0×10^6^ cells and grown at 37°C 5% CO_2_. On day 3, cells were supplemented with 50% original volume with fresh BMDM media. WT and *Trim14*^-/-^ immortalized BMDMs (iBMDMs) were generated with the Cre-J2 virus and weaned off MCSF as described in Nardo et. al^85^. iBMDMs and RAW 264.7 cells were cultured in DMEM, 10% heat inactivated FBS, and 2% HEPES (Cytiva) and grown at 37°C 5% CO_2_. T cells were prepared from whole mouse spleens and or lymph nodes and placed in R10 media (1X RPMI 1640 (Gibco), 10% heat inactivated FBS, 2 mM L-Glutamine). Spleens and lymph nodes were mashed through a 40 µM cell strainer and washed with R10 media. Cells were centrifuged at 1600 rpm for 5 minutes. The remaining cell pellet was resuspended in 1 mL 1X ACK lysis buffer (Quality Biological) for spleens and kept in the dark for 2 min at room temperature. Cells were run through a second 40 µM cell strainer and washed with R10 media. R10 media was immediately added to neutralize the lysis. Splenocytes and/or lymph node cells were centrifuged at 1600 rpm for 5 minutes for downstream assays.

## Method Details

### M. tuberculosis infection

For *M. tuberculosis* infections *ex vivo*, low passage Mtb was prepared by growing it to log phase (OD_600_ 0.6–0.8). Bacterial cultures were spun at 58 rcf for 5 min to remove large clumps. The bacteria were then pelleted at 2,103 rcf for 5 min and washed with 1X PBS. The wash step was repeated twice. The resuspended bacterial cultures were sonicated at 70% amplitude for 10 s and repeated three times (Branson Ultrasonics Corp.) followed by a low-speed spin (58 rcf) to remove remaining clumps. The bacteria were diluted in DMEM (HyClone) + 10% horse serum (Gibco) for *ex vivo* infections or 1X PBS for *in vivo* infections. For *ex vivo* infections, plates containing cells and bacteria were spun at 234 rcf for 10 min to synchronize infection.

### *In vivo* Mtb infections

All Mtb infections were performed using procedures approved by Texas A&M University Institutional Care and Use Committee. The Mtb inoculum was prepared as described above. Age- and sex-matched mice were infected via inhalation exposure using a Madison chamber (Glas-Col) calibrated to introduce ∼100-200 CFUs (low dose) or ∼750 CFUs (medium dose) per mouse. For each infection, approximately 3 mice were euthanized immediately, and their lungs were homogenized and plated to verify the inoculum. Infected mice were housed under BSL3 containment and monitored daily by lab members and veterinary staff. Survival analysis mice were weighed daily and euthanized at 15% body weight loss from starting weight. At the indicated time points, mice were euthanized, and tissue samples were collected. Blood was collected in serum collection tubes, allowed to clot for 1-2 hrs at RT, and spun at 1000 rpm for 10 min to separate serum. Organs were divided for the following readouts: CFUs/protein: inferior lung lobe/middle lobe and 1/2 spleen; histology: superior lung lobe, 1/2 spleen; and flow cytometry: left lung lobe. For histological analysis, organs were fixed for 24 hrs in 10% neutral buffered formalin and moved to ethanol (lung/ heart/ thymus/ lymph nodes/ spleens). Organs were further processed as described below. For CFU enumeration, organs were homogenized in 5 ml PBS + 0.1% Tween-80, and serial dilutions were plated on 7H10 plates. Colonies were counted after plates were incubated at 37°C for 3 weeks.

### Flow cytometry

For TMRE assays examining mitochondrial membrane potential, cells were harvested using 1X PBS-EDTA (Technova). Cell suspensions were spun at 400 rcf for 5 min and resuspended in 1X PBS 4% FBS. Cells were stained for 20 min at 37°C in 25 nM TMRE and washed in 1X PBS 4% FBS before acquisition on a LSR Fortessa X20 (BD Biosciences) with the PE (585/12) channel. For mitochondrial ros analysis, the mitochondria superoxide dye mitoSOX (Thermo Fisher Scientific) was used. Cells were stained with 5 µM mitoSOX for 10 min at 37°C in 1X PBS 2% FBS. Cells were then washed twice with 1X PBS 2% FBS and analyzed by flow cytometry using the PE (585/12) channel.

For apoptosis assays, cells were treated with 10 µM ABT-737 (MedChemExpress) for 6 hours and harvested with 1X PBS-EDTA. Cell suspensions were stained with 5 μg/mL propidium iodide and 25 nM annexin-V (APC, eBioscience) in Annexin binding buffer (Biolegend) for 5 min at room temperature. Cells were washed in 1X PBS 4%FBS and analyzed by flow cytometry using PE (585/12) and APC (670/30) channels. To measure apoptosis in Mtb-infected macrophages, iBMDMs were infected with mCherry Mtb for 6 hours. Cells were harvested with 1X PBS-EDTA and spun at 400 rcf for 5 min. Cells were stained with 400 nM Apotracker Green (Biolegend) for 20 minutes at room temperature. Following staining, cells were washed with 1X PBS 2% FBS twice and fixed in 4% paraformaldehyde (PFA) for 15 min at room temperature. Following fixation, cells were washed with 1X PBS 2% FBS twice and stored at 4°C prior to acquisition on the LSR Fortessa X20.

For macrophage/OT-I splenocytes assays, cells were harvested by PBS lifting and washed with 5% FBS/PBS-EDTA. Cells were stained using the following anti-mouse antibodies in 5% FBS/PBS-EDTA and 10% BD Horizon Brilliant Stain Buffer Plus (BD Biosciences) for 30 minutes at 4°C: CD3 BV650 (Biolegend), CD8 PerCP (Biolegend), CD69 BV421 (Biolegend), CD45 AF532 (Invitrogen). Cells were then washed two times in 1X PBS. Live/dead staining was performed for 20 min in the dark using Ghost dye 510 (Tonbo) at room temperature. Cells were washed two times in 5% FBS/PBS-EDTA before fixing in 4% PFA for 15 min at room temperature. Following fixation, cells were washed with 5% FBS/PBS-EDTA three times and stored at 4°C overnight. Intracellular staining was preformed using a Foxp3 Transcription Factor Staining Permeabilization/Fixation Kit (eBioscience). Briefly, cells were permeabilized in permeabilization buffer for 30 minutes at room temperature. Cells were then stained using the following anti-mouse antibodies in permeabilization buffer and 10% BD Horizon Brilliant Stain Buffer Plus (BD Biosciences) for 30 minutes at 4°C: IFNγ FITC (Biolegend), Ki67 BV605 (Biolegend), and Granzyme B Pacific Blue (Biolegend). Following two washes with permeabilization buffer, samples were resuspended in 5% FBS/PBS-EDTA and analyzed on the Cytek Aurora. See **Fig. S7E** for flow gating strategy.

For analysis of lung cell populations in Mtb-infected mouse lungs, the left lung lobe was harvested and washed with 1X PBS twice. Lungs were minced and digested in digestion buffer (70 µg/mL Liberase [Roche] and 50 µg/mL DNase I [Worthington Biochemical] in RPMI 1640 [HyClone]) for 30 min at 37°C and 5% CO_2_. Single cell suspensions were made by passing homogenates through 70 µM followed by 40 µM cell strainers. Live/dead staining was performed for 20 min in the dark using Ghost dye 510 (Tonbo) at room temperature. Fc receptors were blocked using CD16/CD32 monoclonal antibody (eBioscience). Cells were stained with antibodies against surface markers for innate immune populations: CD45 (Brilliant Violet 785, 30-F11, BioLegend), CD11b (eFluor 450, M1/70, eBioscience), CD11c (Brilliant Violet 605, N418, Biolegend), CD170 (FITC, S17007L, Biolegend), Ly6G (PerCP-Cy5.5, RB6-8CS, eBioscience), Ly6C (APC, HK1.4, eBioscience), CD206 (AF-700, C068C2, Biolegend), and B220 (Alexa Fluor 780, RA3-6B2, eBioscience), and MHC-II (Super Bright 702, M5/114.15.2 eBioscience). Cell were also stained with antibodies against surface markers for adaptive immune populations: CD45 (Brilliant Violet 785, 30-F11, BioLegend), CD3 (Pacific Blue 450, 17A2, Biolegend), CD11c (Brilliant Violet 605, N418, Biolegend), CD62L (FITC, MEL-14, BD), CD44 (PE, IM7, BD), CD8 (PE-eFluor 610, 53-6.7 eBioscience), NK1.1 (APC, PK136, eBioscience), CD4 (Alexa Fluor 700. RM4-5, Biolegend), CD138 (PerCP/Cy5.5, 281-2, Biolegend), and B220 (APC/Cy7, RA3-6B2, Biolegend). Cells were washed two times with 1X PBS 2% FBS before fixing in 4% PFA for 15 min at room temperature. Following fixation, cells were washed with 1X PBS 2% FBS twice and stored at 4°C in the dark prior to acquisition on the LSR Fortessa X20. Total lung cell counts were based on live single cells (Ghost low/−), after doublet exclusion (FSC-H/FSC-A), in 200 µL or 1/3 lung lobe. Forward scatter (FSC)/ Side scatter (SSC) was used to differentiate infiltrating macrophages (CD45^+^ CD11b^+^ Ly6G^−^ CD11c^low/−^ SSC^hi^ FSC^mid^) and monocytes (CD45^+^ CD11b^+^ Ly6G^−^ CD11c^low/−^ SSC^lo^ FSC^mid^) based on cell size and complexity. See **Fig. S6A-B** for flow gating strategies.

Flow cytometry experiments were performed in the Naresh K. Vashisht College of Medicine Flow Cytometry Cell Sorting Facility at Texas A&M Health Science Center and VMC Flow Cytometry Shared Resource at VUMC.

### Histopathology

Mtb-infected mouse lungs were fixed in 10% neutral buffered formalin and stored overnight at 4°C before removal from the BSL-3 facility. Tissues were processed, paraffin embedded, sectioned at 5 µm, and stained with hematoxylin and eosin by AML Laboratories. Lung pathology was assessed in a blinded manner by a board-certified veterinary pathologist. Pulmonary inflammatory burden was quantified from scanned images of one lung lobe per mouse using QuPath Bioimage Analysis software, version 0.7.0. Histiocytic inflammation was measured as the percentage of total lung cross-sectional area occupied by histiocytic inflammatory infiltrates. Additional histopathologic features were quantified using 500 × 500 µm grid overlays. Neutrophilic inflammation was scored as the percentage of grid squares containing neutrophils, whereas neutrophil clusters were defined as grid squares containing more than five closely apposed neutrophils. Necrotic debris was defined as extracellular, circular, densely basophilic nuclear remnants smaller than erythrocytes, measuring less than 6 µm in diameter, and was quantified as the percentage of grid squares containing necrotic debris. For immunohistochemistry (IHC) of Mtb infected lungs, paraffin embedded blocks were processed by the VUMC Translational Pathology Shared Resource and probed for Cleaved Caspase 3 (CST # 9664). QuPath Bioimage analysis was used to quantify and identify CASP3^+^ cells (DAB staining).

### Gene expression analysis by qRT-PCR

For mouse tissue and cell samples, RNA was extracted using the Direct-zol RNAeasy kits (Zymogen) per manufacturer’s protocol. cDNA was synthesized using the Bio-Rad iScript Reverse Transcription Supermix kit. RT-qPCR was performed in technical triplicate using PowerUp SYBR Green Master Mix and analyzed on a QuantStudio 6 Real-Time PCR System (Applied Biosystems). Gene expression changes were normalized to *Actb* levels. qRT-PCR primer sequences can be found in **Table 1**.

### mRNA sequencing

Bulk RNA-seq was performed in biological triplicate by the Texas A&M AgriLife Genomics and Bioinformatics Service Core using the Illumina NovaSeq 6000 platform. Reads were aligned to the *Mus musculus* reference genome build GRCm39 and analyzed using ROSALIND software (San Diego, CA), which uses a hyperscale architecture developed by ROSALIND, Inc. Differentially expressed genes were identified using a p-value threshold of <0.05. Volcano plots were generated in GraphPad Prism. Bulk RNA-seq data have been deposited in GEO under accession number GSE336086.

### Seahorse metabolic analysis

BMDMs were seeded in Agilent Seahorse cell culture microplates at 5 ×10^4^ cells/well in 96-well plates or 2×10^5^ cells/well in 24-well plates. The following day, cells were stimulated with 10 ng/mL LPS for 4 h or overnight and treated with 10 µM ABT-737 for 6 h. For Mtb infection experiments, BMDMs were infected as described above at the indicated multiplicities of infection for 24 h. The Seahorse XF Mito Stress Test Kit and sensor cartridge were prepared according to the manufacturer’s instructions, with modification of the first injection to include glucose, as previously described by Van den Bossche et al.^86^ and Weindel et al.^27^. Normalization was performed using bicinchoninic acid assay absorbance values (ThermoFisher Scientific) and total cell counts determined by DAPI staining on a Lionheart plate reader (Agilent).

### Plate based assays for determination of cell death

Cells were plated in 96-well half area black clear bottom plates at 2.5 ×10^4^ cells/well in 100 µl of media. The following day, inhibitors were added to the well 1 hr prior to start of experiment as follows: 10 µM Q-VD-OPH (MedChemExpress), 5 µM Stattic (MedChemExpress), and 5 µM CsA (MedChemExpress). To induce the NLRP3 and AIM2 inflammasomes, cells were stimulated with 10 ng/mL LPS (Invivogen) for 3 hrs. To analyze the NLRP3 inflammasome, cells were treated post-LPS with 5 mM ATP (ThermoFisher Scientific). For AIM2 inflammasome treatment, cells were transfected post-LPS with 1 μg/mL poly dA:dT (Invivogen) transfection with lipofectamine (ThermoFisher Scientific) at a 3:1 ratio. For necroptosis induction, cells were activated with 100 ng/mL recombinant murine TNFα (PeproTech), 500 nM SMAC mimetic birinapant (MedChemExpress), and 20 µM pan-caspase inhibitor z-VAD-FMK (MedChemExpress). To analyze apoptosis, cells were treated with 10 µM ABT-737 (MedChemExpress), 100 nM Staurosporine (MedChemExpress), and 50 µM Etoposide (MedChemExpress). For overexpression cell death using the tetracycline inducible system, cells were treated with doxycycline hyclate (Millipore Sigma) before cell death induction. For Mtb infections, cells were infected with Mtb as described above. 5 ug/ml Propidium iodide (ThermoFisher Scientific) was added with the cell death agonists. Total cell number was quantified by either using NucBlue (ThermoFisher Scientific) in 1x PBS or propidium iodide in 0.1% Triton X to completely permeabilize cells (100% cell death). Live cell imaging was done using 4x magnification on either a Lionheart plate reader or Cytation 5 (Biotek). Propidium iodide florescence was also measured by microplate reader (Tecan Spark) at 535/617 nm. Post-run analysis was conducted using Gen5 version 3.15 (Biotek) software.

### Protein analysis by immunoblot

Cells were seeded in 12-well plates at 5 ×10^5^ cells/well. Cell lysates were lysed in either 1% SDS lysis buffer or 1X RIPA buffer. Protein samples separated by SDS-PAGE and transferred to 0.45 µM PVDF-FL membranes (Millipore-Sigma). Membranes were blocked for 1h at room temperature in 5% BSA in 1X TBS. Blots were incubated overnight at 4°C with the following antibodies: Actin (Abcam ab6276, 1:5000), Cleaved Casp3 (CST 9664, 1:1000), Cytochrome C (Abcam ab133504, 1:1000), Flag M2 (Sigma-Aldrich F3165, 1:1000), mCherry (CST 43590, 1:1000), Bcl-xl (CST 2764, 1:1000), Bax (CST 2772, 1:1000), Stat3 (CST 9139, 1:1000), Ser754 Stat3 (CST 98543, 1:1000), Ndufb8 (CST 73951, 1:1000), Tom20 (Sigma-Aldrich MABT166, 1:1000), Vdac1 (Biolegend MMS-5205, 1:1000), and Trim14 (Invitrogen PA5-50806, 1:500). Three 5 min washes were done with 1X TBST (0.1% Tween-20) between each incubation. Membranes were incubated with appropriate LiCOR fluorescent secondary antibodies (1:5000) in 5% BSA/TBS for 2h at RT prior to imaging on a LiCOR Odyssey Fc Imager.

### Mitochondria isolation by biochemical fractionation

Cells were seeded at 5 ×10^6^ cells in 6 cm dishes or 1×10^7^ cells in 10 cm dishes. Cells were lifted with 1x PBS-EDTA and pelleted at 600 rcf for 5 min at 4°C. The cells were resuspended in 500 µL 1X Cytosolic extraction Buffer (Cyto Buffer) (250 mM sucrose, 10 mM Tris (pH 7.4), and 1X Halt Protease/Phosphatase Inhibitor Cocktail) on ice for 10 min. A portion (1/10) volume was taken for whole cell lysate for immunoblot analysis (WCL). Cells were homogenized using a glass tight-fitting pestle B with 60 passes on ice. Homogenates were spun at 1000 rcf for 10 min at 4°C twice while saving the supernatant after each spin. The remaining supernatant collected was placed in a fresh tube and centrifuged at 8000 rcf for 30 minutes at 4°C. The supernatant was collected into a fresh tube. The remaining mitochondria pellet were used for mito-Flow analysis or lysed in 6x sample buffer with 5 mM DTT for immunoblot analysis.

For mitochondria proteinase k protection assays, the mitochondria pellet was resuspended in 1X Cyto Buffer and divided evenly into two tubes. In one tube, 50 µg/ml of Proteinase K (Roche) was added for 15 minutes on ice. To stop the reaction, 1 mM PMSF was added for 5 min on ice. A 1% triton x-100 with proteinase k sample was prepared as a positive control to show complete digestion of proteins. The remaining supernatant was centrifuged at 15000 rcf for 30 min at 4°C. The remaining mitochondria pellets were lysed in 6x sample buffer with 5 mM DTT for immunoblot analysis.

For flow cytometry analysis of mitochondria (mito-Flow), the mitochondria pellet was fixed in 1.5% PFA for 20 minutes on ice. Fixed mitochondria were centrifuged at 18000 rcf for 10 min at 4°C (all remaining spins were conducted at this speed) and washed with 500 µL 2% BSA in 1X mito-Flow (300 mM sucrose, 10 mM Tris (pH 7.4), 0.5 mM EDTA). Mitochondria were stained with 200 nM MitoView green (Biotium) for 15 min at room temperature. After centrifugation, mitochondria were permeabilized and stained in 2% BSA in 1X mito-Flow buffer with 0.05% Triton X-100 and Stat3- AF 647 antibody (CST 14062, 1:100) for 1 hour at room temperature. Mitochondria were washed once with 2% BSA in 1X mito-Flow buffer and analyzed on the LSR Fortessa X20 with the FITC (530/30) and APC (670/30) channels.

### Generation of *Stat3* knockout cells using CRSPR/Cas9

Guide RNA sequences were generated to target the 2^nd^ exon of *Stat3* using the Broad Institute online tool (https://portals.broadinstitute.org/gppx/crispick/public) and synthesized by IDT. The gRNA primers were annealed/phosphorylated and ligated into LentiGuide-Puro (Addgene, plasmid 52963). Using lentiviral packaging with PAX2 and VSVG, Flag-Cas9 expressing Raw 264.7 macrophages (Addgene, plasmid 52962) were transduced and selected for using 5 ug/ml puromycin (Invivogen) and 10 ug/ml blasticidin (Invivogen). Knockout efficiency was analyzed by TIDE analysis of a 500 bp region of exon 2. Cells were clonally selected and knockout efficiency was evaluated by immunoblot.

### Tetracycline inducible and stable overexpression cell line generation

To generate tet-iBMDMs and tet-Raw cells, pLenti CMV rtTA3 Blast (Addgene w756-1) was stably transduced using with pLenti CMV Puro DEST (Addgene w118-1) containing 3xFLAG-GFP, mCherry, 3xFLAG-STAT3, 3xFLAG-MTS-STAT3, or 3xFLAG-TRIM14. After 2 days following transduction, cells were selected with 5 ug/ml puromycin and 10 ug/ml blasticidin. To induce expression, doxycycline hyclate (Millipore Sigma) was used as indicated in the figure legends. Stable overexpression cell lines were constructed by transducing cells with pLenti PGK Puro DEST (w529-2) (Addgene 19068).

Site-directed mutagenesis was used to generate 3xFLAG-STAT3 and 3xFLAG-MTS-STAT3 phosphomutants. NEBaseChanger was used to make primers directed to Stat3 at serine 754 to Alanine (A) (phospho-dead) or Glutamic acid (E) (phospho-mimetic). Q5 Hot Start High fidelity DNA polymerase (NEB) was used followed by DpnI (NEB) reaction at 37°C for 3.5 hr. Digested PCR product was transformed into DH5α cells. 3xFLAG-MTS-STAT3 phosphomutants were cloned using NEBuilder HiFi DNA Assembly Reaction Protocol (NEB).

### Determination of mPTP opening by Calcein AM-CoCl_2_ assay

Cells were seeded at 2.5 ×10^5^ cells in 24 well plates. To inhibit mPTP opening, cells were treated with 0.5 µM cyclosporin A (CsA) for 16 hrs. Cells were lifted with 1x PBS-EDTA and pelleted at 400 rcf for 5 min in v-bottom 96 well plates. Cells were washed once with 1x HBSS containing calcium/ magnesium (Hyclone) supplemented with 10 mM Hepes and 2 mM L-glutamine. Next, cells were stained with 0.5 µM calcein AM (ThermoFisher Scientific) and 2 mM CoCl_2_ (Sigma-Aldrich) for 15 minutes at 37°C. Following one wash with 1x HBSS, samples were analyzed on the LSR Fortessa X20 with the FITC channel (530/30).

For Mtb-infected cells, cells were seeded in 96-well half-area black, clear-bottom plate at 2.5 ×10^4^ cells/well in 100 µL of medium. Calcein AM–CoCl₂ staining was performed as described above, except fluorescence was measured using a Tecan Spark plate reader at 485/535 nm excitation/emission.

### Macrophage/OT-I splenocyte co-culture Mtb-OVA infections

Cells were seeded in 24-well plates at 2.0 ×10^5^ cells/well and infected with Mtb-OVA at a MOI=5 for 8 hours as described above. Mtb-OVA was verified to express CFP-10 fused to OVA peptides sequences (257-264 (OT-I) and 323-339 (OT-II)) by PCR as described in Einarsdottir et. al^87^ (**Fig. S7C**). Mtb gDNA was prepared using the CTAB method as described in^88–90^. CFP-10 control primers:CACCTCTAGAGCTCGCGCAGGAGCGTGAAGAAG (sense, CFP10 5’) and TATACATATGGAAGCCCATTTGCGAGGACAGCG (antisense, CFP10 3’) approximately 450 bp (**Fig. S7C**). CFP-10 fused to OVA fragments primers: CFP10 5’ and GGTGGAATTCGCGGCCGGCCTCGTT (antisense, OT-II 3’) approximately 537 bp (**Fig. S7C**). The following PCR reaction with DreamTaq DNA Polymerase (Thermo Fisher Scientific) was used: 95°C 1min, 95°C 30 sec, 68°C 30 sec, 72°C 30 sec, step 2 repeated 20 times, and extension time of 5 min 72°C. At 8 hours post-infection, OT-1 splenocytes were isolated as described above and placed onto Mtb-infected macrophages at a ratio of 1:1 for 16 hrs at 37°C 5% CO_2_. Positive control OT-I splenocytes were treated with 10 nM OVA_257-264_ peptide (MedChemExpress) for 16hrs.

### Statistical analysis

All data are representative of at least two independent experiments with n ≥ 3. For all quantifications, n denotes the number of biological replicates, defined as either individual wells containing cells or individual mice. Data are presented as mean ± SEM. For *in vitro* assays, statistical significance was determined using either a two-tailed Student’s unpaired *t* test, one-way ANOVA followed by Tukey’s test, or two-way ANOVA with Tukey’s *post hoc* test. For *in vivo* mouse infection experiments, statistical significance was determined using the Mann–Whitney U test, based on the assumption that mouse-derived samples were not normally distributed. Survival differences were analyzed using the log-rank Mantel–Cox test. The statistical test used for each experiment is indicated in the corresponding figure legend. All statistical analyses were performed using GraphPad Prism software, version 10, and significance is reported as P values. The threshold for significance was determined by a *P* value of < 0.05, and annotated as \**P* < 0.05, \*\**P* < 0.01, \*\*\**P* < 0.001, and \*\*\*\**P* < 0.0001. For *in vivo* mouse infection experiments, sample size was estimated based on the ability to detect a 0.7 x 10^10^ CFU difference between groups, assuming a standard deviation of 0.3 x 10^10^ CFU for each population. Using a two-sided α of 0.05 and 80% power, we calculated that a minimum of five age- and sex-matched mice per group per time point was required to detect a statistically significant difference by t test. Accordingly, at least five mice per genotype per time point were used to assess infection-related outcomes.

