## Supplemental Figs for "Mitochondrial STAT3-mediated suppression of apoptosis constrains antimycobacterial immunity"

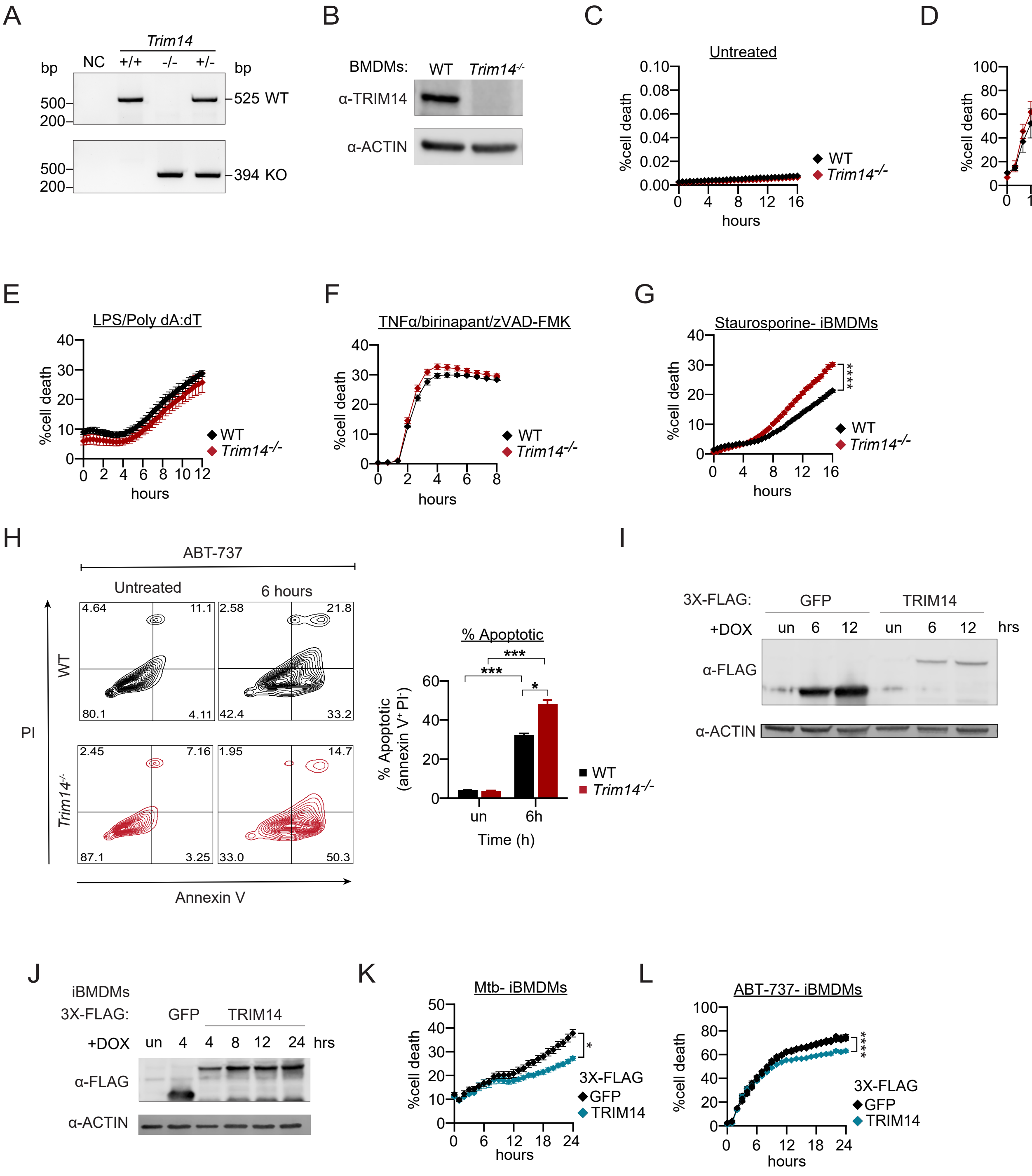
A
Trim14
bp
NC
+/+
-/-
+/-
bp
500
200
525 WT
394 KO
B
BMDMs:
WT
Trim14-/-
α-TRIM14
α-ACTIN
C
Untreated
%cell death
hours
WT
Trim14-/-
D
LPS/ ATP
E
LPS/Poly dA:dT
F
TNFα/birinapant/zVAD-FMK
G
Staurosporine- iBMDMs
****
H
ABT-737
Untreated
6 hours
WT
Trim14-/-
PI
Annexin V
4.64
11.1
80.1
4.11
2.58
21.8
42.4
33.2
2.45
7.16
87.1
3.25
1.95
14.7
33.0
50.3
% Apoptotic
% Apoptotic (annexin V+ PI-)
***
***
*
un
6h
Time (h)
I
3X-FLAG:
GFP
TRIM14
+DOX
un
6
12
hrs
α-FLAG
α-ACTIN
J
iBMDMs
3X-FLAG:
GFP
TRIM14
+DOX
un
4
8
12
24
hrs
α-FLAG
α-ACTIN
K
Mtb- iBMDMs
3X-FLAG
GFP
TRIM14
L
ABT-737- iBMDMs
****

Figure S1

**Supplemental Figure Legends:**

**Figure S1. Trim14 restrains macrophage apoptotic sensitivity**

A) Representative PCR genotyping gel confirming WT, *Trim14*$^{+/-}$, and *Trim14*$^{-/-}$ genotypes. Genotyping PCR details are provided in the Methods.

B) Immunoblot of Trim14 protein levels in WT and *Trim14*$^{-/-}$ BMDMs. Actin was used as loading control.

C) Cell death in untreated WT and *Trim14*$^{-/-}$ BMDMs.

D) Cell death in NLRP3 inflammasome activated (10 ng/mL LPS and then 5 mM ATP) in WT and *Trim14*$^{-/-}$ BMDMs.

E) Cell death in AIM2 inflammasome activated (10 ng/mL LPS and then transfection of 1 μg/mL poly dA:dT ) in WT and *Trim14*$^{-/-}$ BMDMs.

F) Cell death in necroptosis induced (100 ng/mL TNFα, 500 nM birinapant, and 20 μM pan-caspase inhibitor z-VAD-FMK) in WT and *Trim14*$^{-/-}$ BMDMs.

G) Cell death in staurosporine treated (0.1 μM) WT and *Trim14*$^{-/-}$ iBMDMs.

H) Flow cytometry analysis of annexin-V in WT and *Trim14*$^{-/-}$ iBMDMs treated with 10 μM ABT-737 for 6hr. Quantification of % apoptotic (right) (% annexin V + PI -).

I) Immunoblot analysis tetracycline inducible overexpression Raw 264.7 macrophages (treated with 333ng/ml doxycycline for 6hr and 12hr) expressing 3X-Flag GFP or 3X-Flag TRIM14. Actin was used as a loading control.

J) As in (L) but in tetracycline inducible overexpression iBMDMs (4, 8, 12, 24 hr treated with 333ng/ml doxycycline).

K) Cell death in Mtb infected (MOI=5) tetracycline inducible overexpression iBMDMs (4hr pretreatment with 333ng/ml doxycycline) expressing 3X-Flag GFP or 3X-Flag TRIM14.

L) As in (K) but treated with 10 μM ABT-737.

Statistical analysis: *p < 0.05, **p < 0.01, ***p < 0.001, ****p < 0.0001. Statistical differences were determined for (C-H, K-L) using two-way ANOVA with Tukey's post-test.

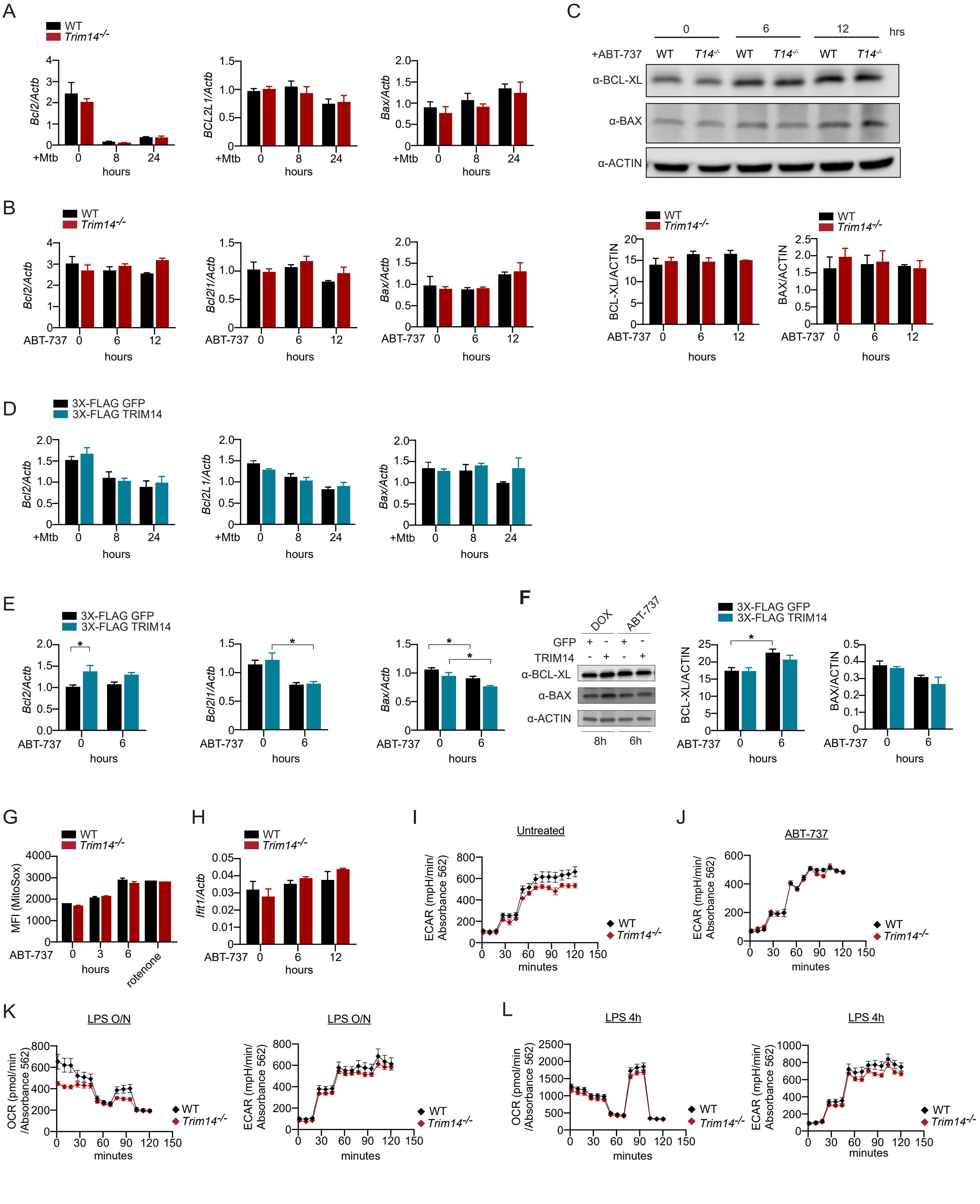
A
WT
Trim14-/-
Bcl2/Actb
BCL2L1/Actb
Bax/Actb
+Mtb
hours
C
+ABT-737
0
6
12
hrs
WT
T14-/-
α-BCL-XL
α-BAX
α-ACTIN
B
Bcl2/Actb
Bcl2l1/Actb
Bax/Actb
ABT-737
BCL-XL/ACTIN
BAX/ACTIN
D
3X-FLAG GFP
3X-FLAG TRIM14
E
F
DOX
ABT-737
GFP
TRIM14
8h
6h
G
MFI (MitoSox)
rotenone
H
Ifit1/Actb
I
Untreated
ECAR (mpH/min/ Absorbance 562)
minutes
J
ABT-737
K
LPS O/N
OCR (pmol/min /Absorbance 562)
L
LPS 4h

Figure S2

**Figure S2. Trim14 does not broadly alter apoptotic gene expression during Mtb infection.**

A) qRT-PCR gene expression of *Bcl2*, *Bcl2l1*, and *Bax* in uninfected and Mtb-infected (MOI=5 8hr and 24hr) WT and *Trim14*$^{-/-}$ BMDMs. Gene expression was normalized to *Actin*.

B) As in (A) but with ABT-737 treatment (10 μM 6hr and 12hr).

C) Immunoblot analysis of Bcl-xl and Bax in untreated and ABT-737 treated (10 μM 6hr and 12hr) WT and *Trim14*$^{-/-}$ BMDMs (top). Quantification of Bcl-xl and Bax (bottom) was normalized to Actin protein levels.

D) As in (A) but Mtb infection (MOI=5 8hr and 24hr) in tetracycline inducible overexpression iBMDMs (treated with 333ng/ml doxycycline for 8hr pre-infection) expressing 3X-Flag GFP or 3X-Flag TRIM14.

E) As in (B) but in tetracycline inducible overexpression iBMDMs (treated with 333ng/ml doxycycline for 8hr pre-infection) expressing 3X-Flag GFP or 3X-Flag TRIM14.

F) As in (C) but with tetracycline inducible overexpression iBMDMs (treated with 333ng/ml doxycycline for 8hr pre-infection) expressing 3X-Flag GFP or 3X-Flag TRIM14 and then treated with 10 μM ABT-737 for 6hr (left). Quantification of Bcl-xl and Bax (right) was normalized to Actin protein levels.

G) Flow cytometry analysis of MitoSox in untreated and 10 μM ABT-737 treated (3 and 6 hours) WT and *Trim14*$^{-/-}$ iBMDMs. 2.5 μM for 2 hours was used as a ROS inducer.

H) As in (B) but measuring *Ifit1* gene expression.

I) Extracellular acidification rates (ECAR) measured by Agilent Seahorse Metabolic Analyzer in untreated WT and *Trim14*$^{-/-}$ BMDMs.

J) As in (H) but 10 μM ABT-737 treatment for 6hr.

K) Oxygen consumption rate (OCR) (left) and Extracellular acidification rates (ECAR) (right) measured by Agilent Seahorse Metabolic Analyzer in WT and *Trim14*$^{-/-}$ BMDMs: LPS treated (16h 10 ng/ml).

L) As in (J) but treated with LPS (4h 10 ng/ml).

Statistical analysis: *p < 0.05, **p < 0.01, ***p < 0.001, ****p < 0.0001. Statistical differences were determined for (A-H) using two-way ANOVA with Tukey's post-test.

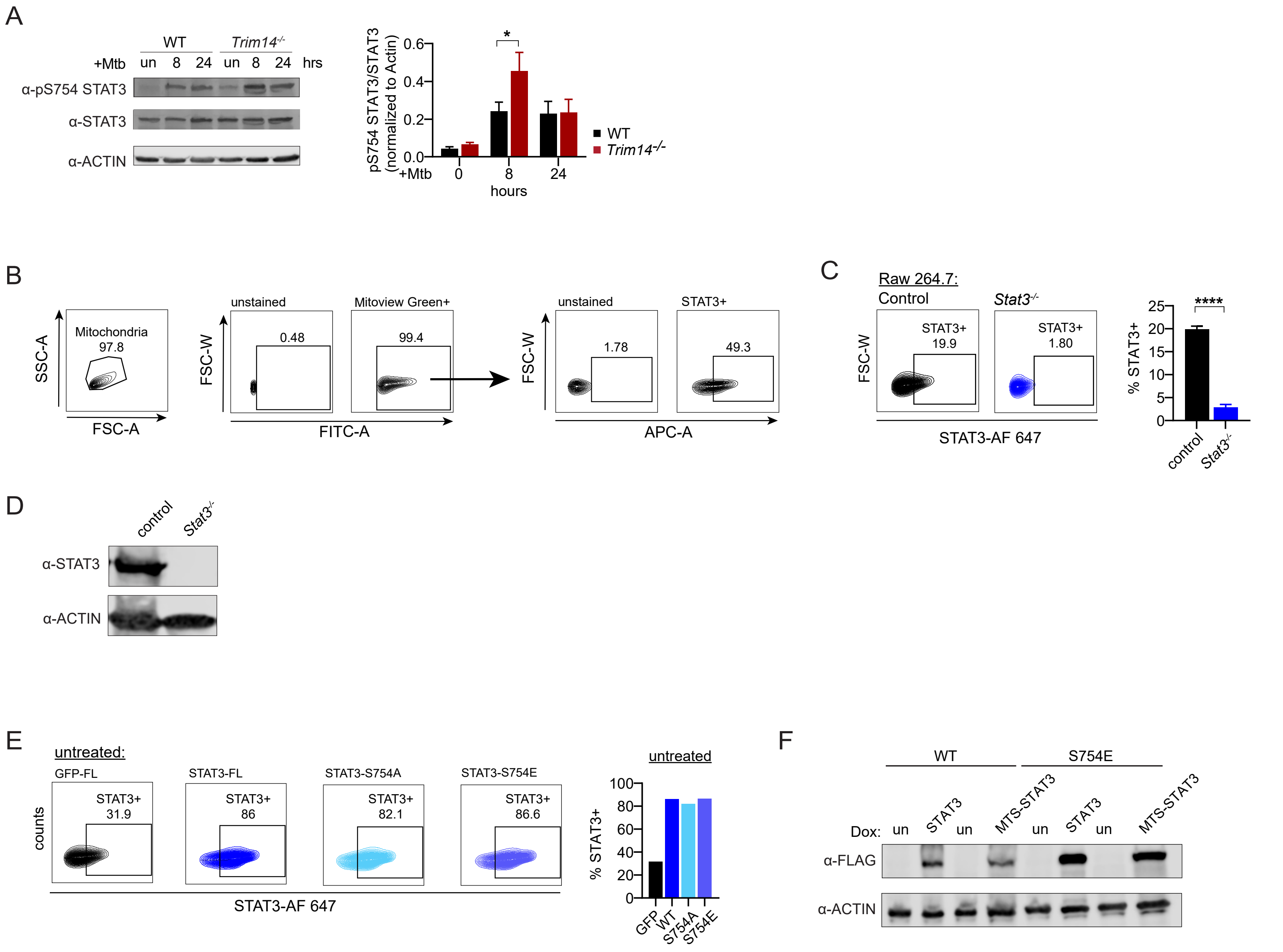
A
WT
Trim14-/-
+Mtb un 8 24 un 8 24 hrs
α-pS754 STAT3
α-STAT3
α-ACTIN
pS754 STAT3/STAT3 (normalized to Actin)
+Mtb 0 8 24
hours
WT
Trim14-/-
B
Mitochondria 97.8
SSC-A
FSC-A
unstained
Mitoview Green+
0.48
99.4
FSC-W
FITC-A
unstained
STAT3+
1.78
49.3
APC-A
C
Raw 264.7:
Control
Stat3-/-
STAT3+ 19.9
STAT3+ 1.80
STAT3-AF 647
% STAT3+
control
Stat3-/-
D
α-STAT3
α-ACTIN
E
untreated:
GFP-FL
STAT3-FL
STAT3-S754A
STAT3-S754E
STAT3+ 31.9
STAT3+ 86
STAT3+ 82.1
STAT3+ 86.6
counts
STAT3-AF 647
GFP WT S754A S754E
F
WT
S754E
Dox: un STAT3 un MTS-STAT3 un STAT3 un MTS-STAT3
α-FLAG
α-ACTIN

Figure S3

**Figure S3. Mito-flow validation and analysis of Trim14-dependent mitochondrial Stat3 localization**

A) Immunoblot of pS754 Stat3 and Stat3 levels in WT and *Trim14*$^{-/-}$ BMDMs infected with Mtb (MOI=5) for 8h and 24h. Actin was used as a loading control.

B) Mito-flow gating strategy. Enriched mitochondria were gated on Mitoview Green positivity.

C) Mito-flow analysis of Stat3 mitochondrial association in untreated control and *Stat3*$^{-/-}$ Raw 264.7 macrophages. Quantification on the right.

D) Immunoblot analysis of Stat3 levels in control and *Stat3*$^{-/-}$ Raw 264.7 macrophages.

E) Mitoflow analysis of untreated stably expressing iBMDMs expressing 3X-Flag GFP, 3X-Flag STAT3, 3X-Flag STAT3 S754A, or 3X-Flag STAT3 S754E. Quantification on right. n=1

F) Immunoblot of Flag in tetracycline-inducible iBMDMs expressing 3X-Flag-STAT3, 3X-Flag-STAT3 S754E, or the corresponding MTS-tagged constructs following doxycycline induction (1 µg/mL, 8h). Actin was used as a loading control.

Statistical analysis: *p < 0.05, **p < 0.01, ***p < 0.001, ****p < 0.0001. Statistical differences were determined for (A) using two-way ANOVA with Tukey's post-test and (C) using unpaired two-tailed Student's t test.

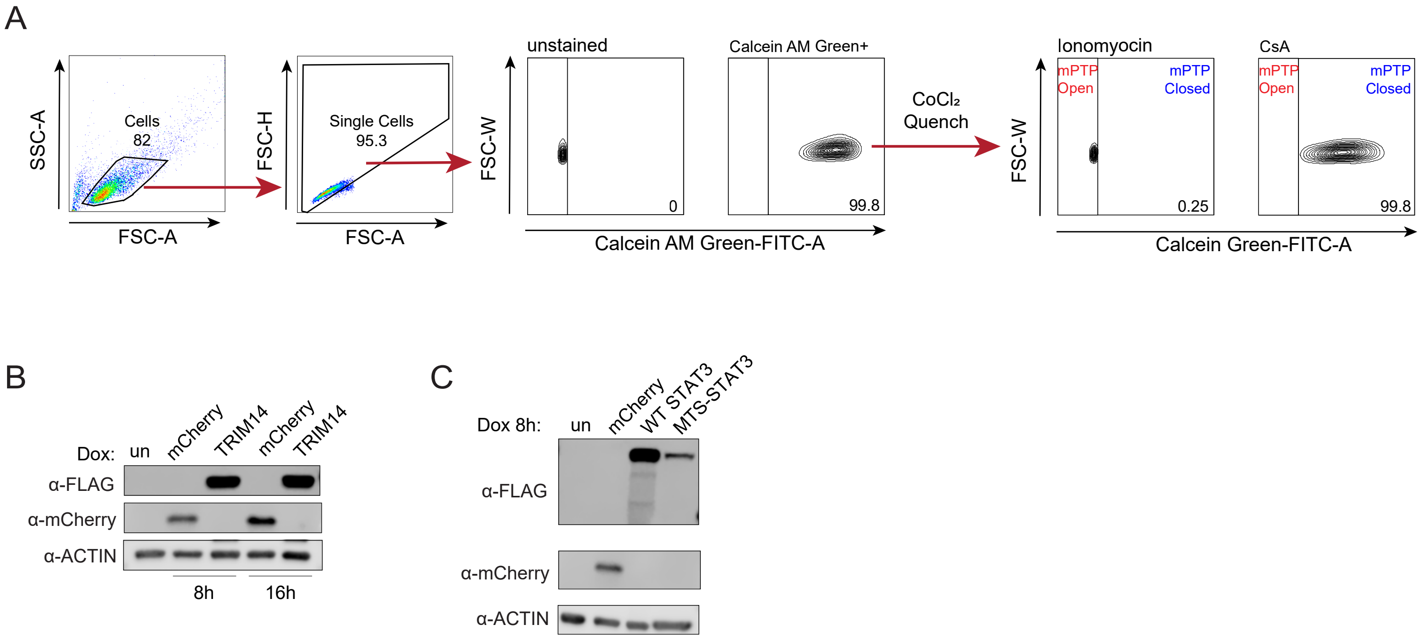
A
SSC-A
FSC-A
Cells
82
FSC-H
Single Cells
95.3
FSC-W
unstained
0
Calcein AM Green+
99.8
Calcein AM Green-FITC-A
CoCl₂
Quench
Ionomyocin
mPTP
Open
mPTP
Closed
0.25
CsA
99.8
Calcein Green-FITC-A
B
Dox:
un
mCherry
TRIM14
α-FLAG
α-mCherry
α-ACTIN
8h
16h
C
Dox 8h:
WT STAT3
MTS-STAT3

Figure S4

**Figure S4. Flow cytometric analysis of mPTP opening and immunoblot validation of Trim14 and Stat3 expression.**

A) Flow cytometry gating strategy for calcein fluorescence measured before and after $CoCl_2$ quench.

B) Immunoblot analysis mCherry and Flag in tetracycline inducible overexpression iBMDMs (333ng/ml doxycycline for 8hr or 16hr) expressing mCherry or 3X-Flag TRIM14. Actin was used as a loading control.

C) As in (B) but in tetracycline inducible overexpression iBMDMs (333ng/ml doxycycline for 8hr) expressing mCherry, 3X-Flag STAT3, or 3X-Flag MTS-STAT3.

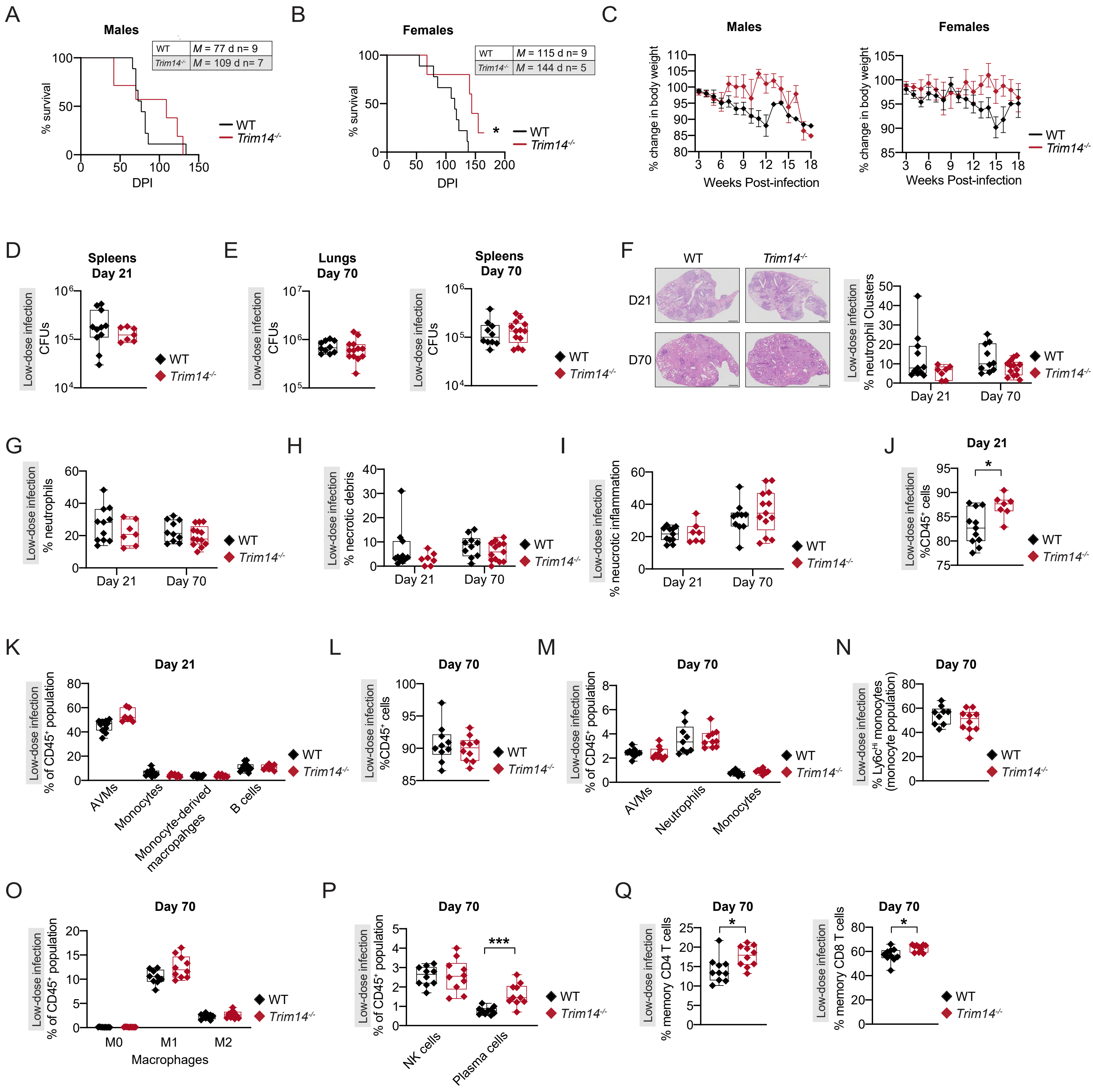

A
Males
WT M = 77 d n= 9
Trim14-/- M = 109 d n= 7
% survival
DPI
WT
Trim14-/-
B
Females
WT M = 115 d n= 9
Trim14-/- M = 144 d n= 5
*
C
Males
Females
% change in body weight
Weeks Post-infection
D
Spleens Day 21
E
Lungs Day 70
Spleens Day 70
Low-dose infection
CFUs
F
D21
D70
% neutrophil Clusters
Day 21
Day 70
G
% neutrophils
H
% necrotic debris
I
% neucrotic inflammation
J
%CD45+ cells
K
% of CD45+ population
AVMs
Monocytes
Monocyte-derived macropahges
B cells
L
M
Neutrophils
N
% Ly6cHi monocytes (monocyte population)
O
M0
M1
M2
Macrophages
P
NK cells
Plasma cells
***
Q
% memory CD4 T cells
% memory CD8 T cells


Figure S5

**Figure S5. Loss of Trim14 reshapes pulmonary immune responses during Mtb infection**

A) Survival analysis of WT (n=9) and *Trim14*$^{-/-}$ (n=7) male mice infected with ~750 Mtb.

B) Survival analysis of WT (n=9) and *Trim14*$^{-/-}$ (n=5) female mice infected with ~750 Mtb.

C) Percent body weight change of WT and *Trim14*$^{-/-}$ mice infected with ~750 Mtb bacilli separated by sex.

D) Spleen CFUs of WT and *Trim14*$^{-/-}$ mice at day 21.

E) Lung and Spleen CFUs of WT and *Trim14*$^{-/-}$ mice at day 70.

F) H&E representative images of WT and *Trim14*$^{-/-}$ mice lungs at day 21 and 70. Scale bar = 800 μM. H&E analysis of neutrophil clusters in WT and *Trim14*$^{-/-}$ mice lungs.

G) As in (F) but neutrophils.

H) As in (F) but necrotic debris.

I) As in (F) but neucrotic inflammation.

J) FACs analysis of CD45$^{+}$ cells in WT and *Trim14*$^{-/-}$ mice lungs at day 21.

K) FACs analysis of AVMs, monocytes, monocyte-derived macrophages, and B cells as a % of CD45$^{+}$ cells in WT and *Trim14*$^{-/-}$ mice lungs at day 21.

L) As in (J) but day 70.

M) FACs analysis of alveolar macrophages, neutrophils, and monocytes as a % of CD45$^{+}$ cells in WT and *Trim14*$^{-/-}$ mice lungs at day 70.

N) FACs analysis of Ly6c$^{Hi}$ monocytes as a % of CD45$^{+}$ cells in WT and *Trim14*$^{-/-}$ mice lungs at day 70.

O) FACs analysis of M0, M1, and M2 macrophages as a % of CD45$^{+}$ cells in WT and *Trim14*$^{-/-}$ mice lungs at day 70.

P) As in (O) but NK cells and plasma cells.

Q) FACs analysis of CD4 and CD8 memory T cells as a % of CD45$^{+}$ cells in WT and *Trim14*$^{-/-}$ mice lungs at day 70.

Statistical analysis: *$p < 0.05$, **$p < 0.01$, ***$p < 0.001$, ****$p < 0.0001$. Statistical differences were determined for (A-B) Mantel-Cox log-rank, (C) two-way ANOVA with Tukey's post-test (D-Q) using Mann-Whitney U test.

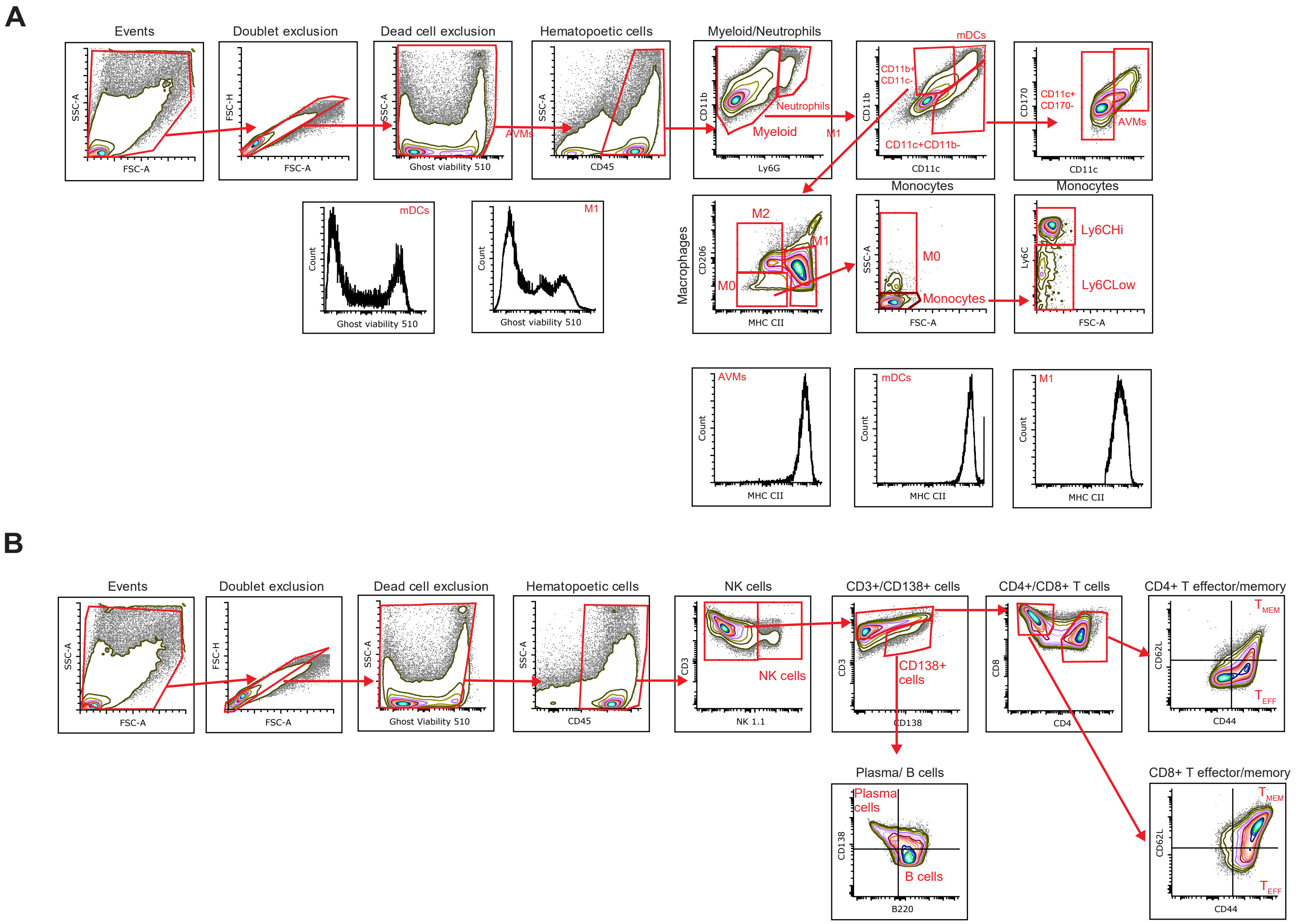
A
Events
Doublet exclusion
Dead cell exclusion
Hematopoetic cells
Myeloid/Neutrophils
mDCs
Neutrophils
Myeloid
AVMs
M1
CD11b+ CD11c-
CD11c+CD11b-
CD11c+ CD170-
Monocytes
M2
M0
Ly6CHi
Ly6CLow
Macrophages
SSC-A
FSC-A
FSC-H
Ghost viability 510
CD45
CD11b
Ly6G
CD11c
CD170
CD206
MHC CII
Ly6C
Count
B
NK cells
CD3+/CD138+ cells
CD4+/CD8+ T cells
CD4+ T effector/memory
CD8+ T effector/memory
Plasma/ B cells
CD138+ cells
Plasma cells
B cells
Ghost Viability 510
CD3
NK 1.1
CD138
CD8
CD4
CD62L
CD44
B220

Figure S6

**Figure S6. Flow cytometry gating strategies for innate and adaptive immune panels.**

A) Gating strategy for innate immune flow cytometry analysis at D21 and D70 post-Mtb infection in WT and *Trim14*$^{-/-}$ mice.

B) As in (A) but adaptive immune flow cytometry panel.

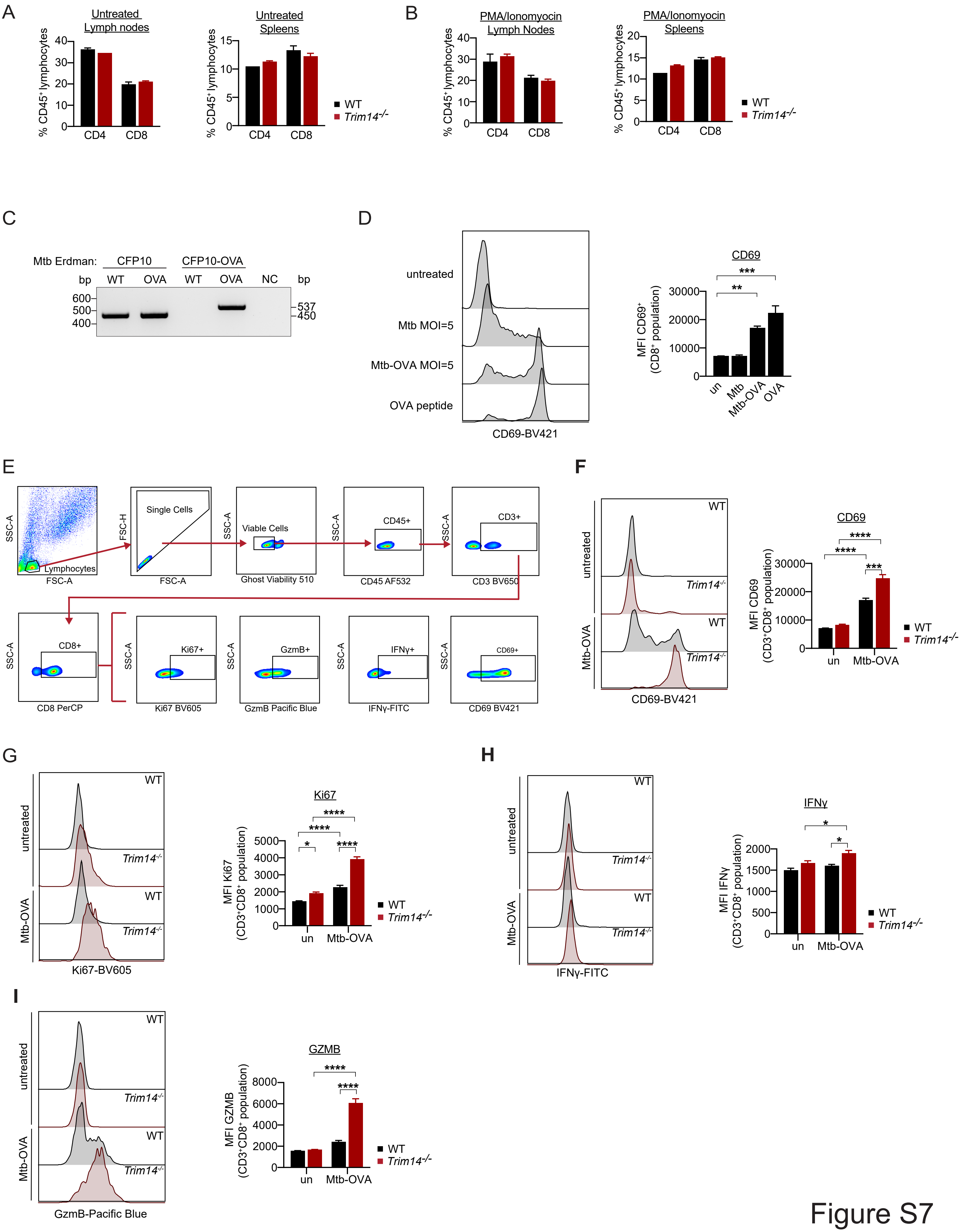
A
Untreated Lymph nodes
Untreated Spleens
% CD45+ lymphocytes
CD4
CD8
WT
Trim14-/-
B
PMA/Ionomyocin Lymph Nodes
PMA/Ionomyocin Spleens
C
Mtb Erdman:
CFP10
CFP10-OVA
bp
WT
OVA
NC
600
500
400
537
450
D
untreated
Mtb MOI=5
Mtb-OVA MOI=5
OVA peptide
CD69-BV421
CD69
MFI CD69+ (CD8+ population)
un
Mtb
Mtb-OVA
OVA
E
SSC-A
FSC-A
FSC-H
Lymphocytes
Single Cells
Viable Cells
Ghost Viability 510
CD45+
CD45 AF532
CD3+
CD3 BV650
CD8+
CD8 PerCP
Ki67+
Ki67 BV605
GzmB+
GzmB Pacific Blue
IFNγ+
IFNγ-FITC
CD69+
CD69 BV421
F
Mtb-OVA
MFI CD69 (CD3+CD8+ population)
G
Ki67-BV605
Ki67
MFI Ki67 (CD3+CD8+ population)
H
IFNγ-FITC
IFNγ
MFI IFNγ (CD3+CD8+ population)
I
GzmB-Pacific Blue
GZMB
MFI GZMB (CD3+CD8+ population)

Figure S7

**Figure S7. *Trim14*$^{-/-}$ macrophages enhance CD8+ T cell activation in a Mtb-OVA co-culture model**

A) FACs analysis of %CD8 and %CD4 surface staining on lymphocytes from lymph nodes and spleens from uninfected WT and *Trim14*$^{-/-}$ mice (n=2 for each genotype).

B) As in (A) but FACs analysis following PMA/Ionomycin stimulation for 4hrs.

C) PCR validation of gDNA from WT Mtb Erdman and Mtb-OVA Erdman strains. CFP-10 was used as a validated secreted virulence factor (~450 bp). CFP-10-OVA was validated in Mtb-OVA (~537 bp).

D) Histogram of CD69 expression from Fig. 7B. MFI quantification on the right.

E) Flow gating strategy for FACs analysis.

F) As in (D) but measuring CD69 expression in uninfected and Mtb-OVA infected WT and *Trim14*$^{-/-}$ iBMDMs.

G) As in (F) but measuring Ki67 expression.

H) As in (F) but measuring IFNγ expression.

I) As in (F) but measuring GZMB expression.

Statistical analysis: *p < 0.05, **p < 0.01, ***p < 0.001, ****p < 0.0001. Statistical differences were determined for (D) using one-way ANOVA with Tukey's post-test, (F-I) two-way ANOVA with Tukey's post-test, and (A-B) using multiple two-tailed Student's unpaired t tests.

**Table 1. qRT-PCR primers.**

| | FWD | REV |
|---|---|---|
| *Bcl2* | CCTGTGGATGACTGAGTACCTG | AGCCAGGAGAAATCAAACAGAGG |
| *Bcl2l1* | GCCACCTATCTGAATGACCACC | AGGAACCAGCGGTTGAAGCGC |
| *Bax* | CCGGCGAATTGGAGATGAACTG | AGCTGCCACCCGGAAGAAGACCT |
| *Ifit1* | CGTAGCCTATCGCCAAGATTTA | AGCTTTAGGGCAAGGAGAAC |
| *Actb* | GGTGTGATGGTGGGAATGG | GCCCTCGTCACCCACATAGGA |
